# Two Axes of White Matter Development

**DOI:** 10.1101/2025.03.19.644049

**Authors:** Audrey C. Luo, Steven L. Meisler, Valerie J. Sydnor, Aaron Alexander-Bloch, Joëlle Bagautdinova, Deanna M. Barch, Dani S. Bassett, Christos Davatzikos, Alexandre R. Franco, Jeff Goldsmith, Raquel E. Gur, Ruben C. Gur, Fengling Hu, Marc Jaskir, Gregory Kiar, Arielle S. Keller, Bart Larsen, Allyson P. Mackey, Michael P. Milham, David R. Roalf, Golia Shafiei, Russell T. Shinohara, Leah H. Somerville, Sarah M. Weinstein, Jason D. Yeatman, Matthew Cieslak, Ariel Rokem, Theodore D. Satterthwaite

## Abstract

Despite decades of neuroimaging research, how white matter develops along the length of major tracts in humans remains unknown. Here, we identify fundamental patterns of white matter maturation by examining developmental variation along major, long-range cortico-cortical tracts in youth ages 5-23 years using diffusion MRI from three large-scale, cross-sectional datasets (total *N* = 2,716). Across datasets, we delineate two replicable axes of human white matter development. First, we find a deep-to-superficial axis, in which superficial tract regions near the cortical surface exhibit greater age-related change than deep tract regions. Second, we demonstrate that the development of superficial tract regions aligns with the cortical hierarchy defined by the sensorimotor-association axis, with tract ends adjacent to sensorimotor cortices maturing earlier than those adjacent to association cortices. These results reveal developmental variation along tracts that conventional tract-average analyses have previously obscured, challenging the implicit assumption that white matter tracts mature uniformly along their length. Such developmental variation along tracts may have functional implications, including mitigating ephaptic coupling in densely packed deep tract regions and tuning neural synchrony through hierarchical development in superficial tract regions – ultimately refining neural transmission in youth.

## INTRODUCTION

White matter (WM) tracts undergo protracted refinement in youth, supporting communication between spatially distributed cortical regions^1,2^. In cortical gray matter, convergent lines of evidence have shown that development progresses heterochronously along the cortical hierarchy defined by the sensorimotor-association (S-A) axis, with unimodal sensorimotor regions developing earlier and transmodal association regions maturing later^3–7^. However, how development varies along white matter tracts – which may provide insight into how different positions along a tract may serve distinct functional roles – has yet to be systematically characterized. Furthermore, despite the intrinsic relationship between white and gray matter, they are typically studied separately. As a result, little is known about how major WM tracts develop in relation to the cortical regions they connect. Here, we sought to define spatial patterns of development in cortico-cortical WM tracts in youth.

Despite decades of research, there is limited understanding of how WM develops along the length of individual tracts. Foundational studies using diffusion MRI established that widespread decreases in mean diffusivity and increases in fractional anisotropy occur in childhood and adolescence^2,8–10^, corresponding to increases in myelination, refinements in axonal caliber, and changes in glia^11^. Notably, nearly all prior work of *in vivo* WM development has averaged measures of microstructure along the entire tract, a methodological approach that implicitly assumes that WM develops synchronously along its length. This approach precludes more spatially precise investigations of developmental variation along tracts and neglects important nuances suggested by animal and post-mortem studies. In animal models, myelination has been shown to vary along axons, which can profoundly impact the velocity and synchrony of neural transmission^12,13^. Infant post-mortem studies examining myelin at discrete WM sites have described broad regional patterns of myelination – e.g., from posterior-to-anterior, inferior-to-superior, and central-to-peripheral WM regions^14,15^. Such regional variation in myelin content and formation warrants investigating how WM development varies continuously along the length of major tracts and identifying the organizing principles underlying this variation.

Examining developmental variation along WM tracts has the potential to provide insights into how tracts mature in relation to the cortices they connect. Understanding this relationship is particularly important given that the primary role of WM tracts is to facilitate communication between distant cortical regions^16^. Long-range association tracts often connect regions that span the S-A axis, a pattern of cortical organization that aligns with hierarchies of cortical anatomy, function, and evolutionary expansion that spans from sensorimotor (lowest ranks) to association (highest ranks) cortices^3^. For example, the inferior fronto-occipital fasciculus connects lower-order visual regions, which largely mature in childhood, to higher-order frontal association regions, which continue developing into early adulthood^6^. Given that cortical activity can lead to preferential myelination of axons^17^, hierarchical cortical development may dynamically interact with WM development. Thus, tracts connecting cortical regions at different positions along the S-A axis may exhibit divergent development near these differing endpoints. Given the growing evidence that sensorimotor cortical regions develop before association regions^4,5,18^ and the plausibility of divergent development along a single tract, prior reliance on whole-tract averages may have masked important patterns of differential development linked to cortical endpoints.

Findings from the few small-scale studies that quantified tissue properties along the length of tracts during youth^19–21^ have suggested the possibility of heterochronous development along tracts. While these studies identified differences in microstructural properties and developmental timing along several tracts, they did not explore the development of tracts in relation to their cortical endpoints. Furthermore, the small samples evaluated in prior work may limit generalizability. Addressing gaps in understanding WM development requires studying along tract development in the context of the cortical hierarchy in large samples.

Here, we aimed to delineate fundamental patterns of WM development by examining developmental variation along major cortico-cortical tracts in youth. We hypothesized that development would not be uniform along a WM tract, but would vary in part based on the position of the tract’s endpoints along the cortical hierarchy. To ensure rigor and assess replicability, we analyzed three independently-acquired, large-scale neuroimaging datasets (total *N* = 2,716) of youth ages 5-23 years old. Our findings challenge the assumption of synchronous development along WM tracts implied by conventional tract-average analyses. As described below, we demonstrate that WM development aligns with two distinct axes. First, WM develops along a deep-to-superficial axis within individual, long-range WM tracts, with superficial tract regions adjacent to the cortex exhibiting greater age-related change compared to deep tract regions. Second, development in superficial tract regions aligns with the cortical hierarchy defined by the S-A axis, such that tract ends adjacent to lower-order sensorimotor cortices mature earlier than those adjacent to higher-order association cortices.

## RESULTS

We characterized white matter (WM) development in youth using diffusion MRI from three large-scale, cross-sectional datasets (total *N* = 2,716). We used the Philadelphia Neurodevelopmental Cohort (PNC; *n* = 1,098; ages 8-23 years) as the discovery dataset and Human Connectome Project: Development (HCP-D; *n* = 567; ages 8-22 years) and Healthy Brain Network (HBN; *n* = 1,051; ages 5-22 years) as replication datasets. First, we investigated variability in developmental effects of mean diffusivity at 100 equidistant nodes (numbered 0-99), or spatial locations, along each cortico-cortical tract. Specifically, we quantified differences in developmental effects between deep tract regions (nodes 46-55) and superficial tract regions (nodes 5-9 and 90-94) after trimming the endmost nodes (0-4 and 95-99) to mitigate partial volume effects. Of note, “superficial” here does not refer to U-fibers, but rather to regions of long-range WM tracts that are closer to the cortex. Second, we studied how developmental patterns of superficial tract regions vary according to their respective cortical endpoints. To do so, we mapped tracts to their cortical endpoints by developing a workflow that combines multiple toolkits. Third, we tested whether the development of superficial tract regions varies along the cortical hierarchy defined by the sensorimotor-association (S-A) axis by mapping S-A ranks of cortical endpoints to superficial tract regions.

### White matter development occurs along a deep-to-superficial axis along tracts

We first characterized spatial patterns of developmental change in mean diffusivity along each tract. We focused on mean diffusivity for three reasons. First, mean diffusivity is more sensitive to developmental changes and more robust to the impact of in-scanner motion than other commonly used measures, including fractional anisotropy^8,22,23^. Second, it can be calculated for both single-shell (PNC) and multi-shell (HCP-D, HBN) acquisitions. Third, mean diffusivity is thought to be less sensitive to partial volume effects in WM adjacent to cortex than fractional anisotropy^24,25^. To model developmental changes in mean diffusivity along each tract, we fit generalized additive models at each of the 100 nodes along each tract. This model included age as a smooth term as well as sex and head motion as linear covariates. We quantified the magnitude of age-related change in mean diffusivity for each node by computing the absolute change in adjusted *R*^2^ (Δ*R*^2^adj) between a full model and a reduced model without the age term. Across tracts, mean diffusivity significantly decreased over development in the age window studied at nearly all nodes (*Q*_FDR_ < 0.05 in 95% of nodes in PNC; >99% in HCP-D; 98% of nodes in HBN).

We first evaluated development along callosal tracts (**Figure 1**). Deep WM exhibited the smallest age-related changes; age effects continuously increased in magnitude toward the superficial regions of each tract. The differences were often quite marked. For example, in the motor segment of the corpus callosum (callosum motor), age explained less than 5% of the variance in mean diffusivity of deep tract regions, but explained approximately 36% of variance in superficial tract regions. To statistically evaluate differences in age effects between deep and superficial tract regions, we adapted a recently-introduced permutation-based enrichment test that accounts for autocorrelated data structures (see Methods for details). Across datasets, we found that age effects were significantly enriched in superficial as compared to deep tract regions in all callosal tracts in all datasets (*Q*_FDR_ < 0.05), with the exception of callosum occipital in HCP-D.

**Figure 1.**
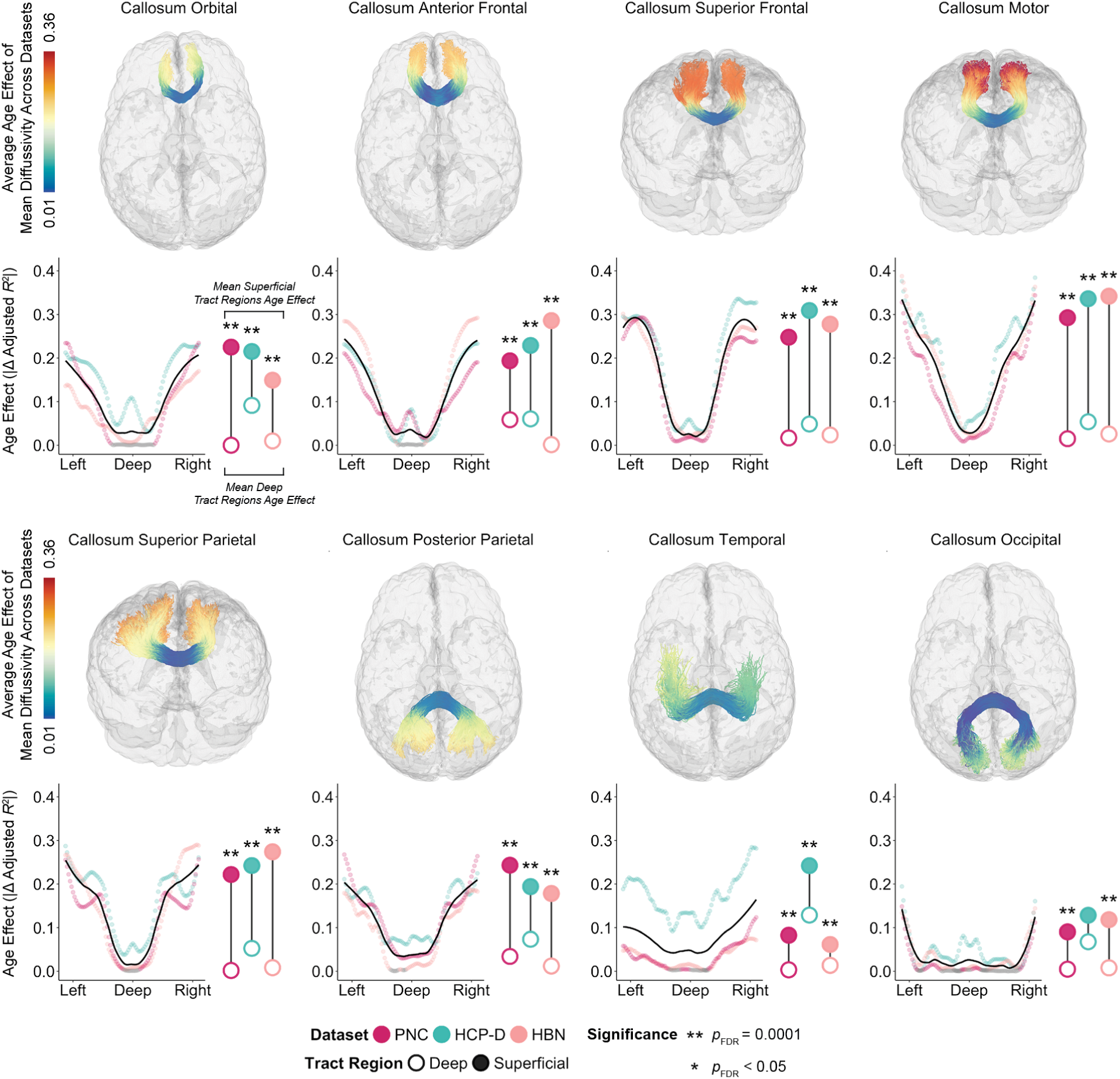
White matter development occurs along a deep-to-superficial axis in callosal tracts. The magnitude of the mean diffusivity age effect varies continuously along each of eight tracts that comprise the corpus callosum. Each callosal tract is shown in a glass brain of an exemplar participant, with that participant’s streamlines colored by the average magnitude of the age effect across datasets. Below each glass brain, we display the magnitude of the age effect at 100 equidistant nodes that span the length of each tract. Data points are colored by dataset: the Philadelphia Neurodevelopmental Cohort (PNC; magenta), Human Connectome Project: Development (HCP-D; teal), and Healthy Brain Network (HBN; coral). Nodes without a significant age association are colored in gray (*Q*_FDR_ > 0.05). The black LOESS-smoothed line shows the overall trend for each tract, averaged across datasets. To the right of each age effect plot, average age effects for superficial (filled circle) and deep (open circle) tract regions are shown for each dataset. Significant differences between age effects of superficial and deep tract regions were assessed using a network enrichment significance test. Stars denote significance levels following FDR correction.

We investigated whether this deep-to-superficial patterning in callosal tracts could generalize to measures derived from more advanced models of diffusion: neurite orientation dispersion and density imaging (NODDI) and mean apparent propagator (MAP) MRI. These models leverage multiple *b*-values (or “shells”) in the dMRI acquisition to characterize features of the cellular environment^22,26–28^. NODDI is a biophysical model that estimates the directional distribution of neurites and compartmental volume fractions^29^. In contrast, MAP-MRI is a signal-based model, which models water molecule displacement without *a priori* assumptions about the tissue environment and estimates properties of the mean apparent propagator directly^30,31^. Using multi-shell diffusion imaging data from HCP-D and HBN, we examined the distribution of age effects of intracellular volume fraction (ICVF; a marker of neurite density) computed from NODDI and return-to-origin probability (RTOP; the probability of water molecules undergoing no net displacement) estimated from MAP-MRI. As with mean diffusivity, we observed a deep-to-superficial patterning of age effects in both ICVF and RTOP **(Figure 2)**, suggesting this gradient is robust across multiple diffusion MRI models.

**Figure 2.**
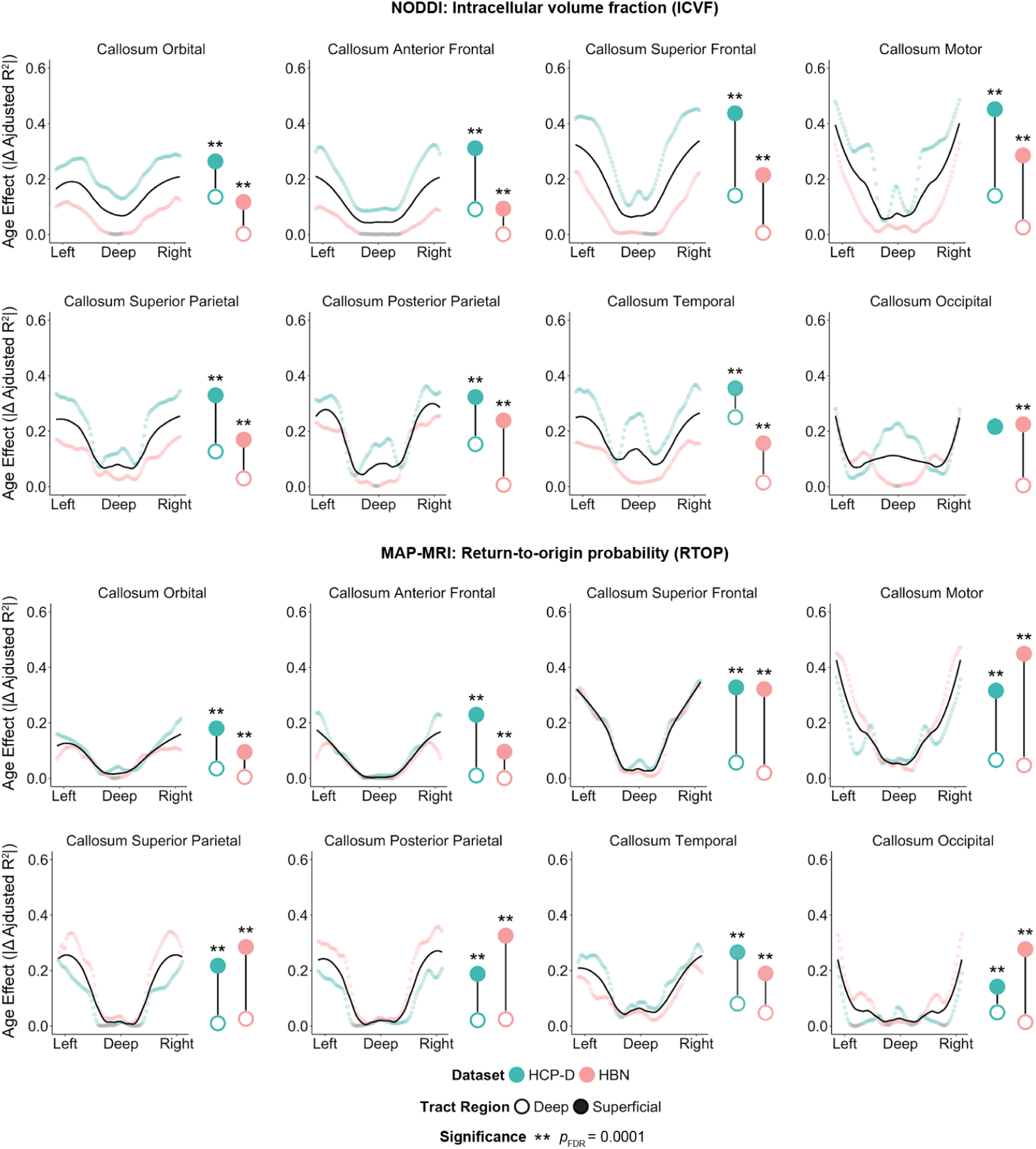
Development of microstructural measures from multi-shell diffusion models occurs along a deep-to-superficial axis in callosal tracts. The magnitudes of the intracellular volume fraction (ICVF; computed from NODDI) and return-to-origin probability (RTOP; computed from MAP-MRI) age effect vary continuously along each of eight tracts that comprise the corpus callosum. We display the magnitude of the age effect for each diffusion measure at 100 equidistant nodes that span the length of each tract. Data points are colored by each dataset with multi-shell diffusion imaging data: Human Connectome Project: Development (HCP-D; teal) and Healthy Brain Network (HBN; coral). Nodes without a significant age association are colored in gray (*Q*_FDR_ > 0.05). The black LOESS-smoothed line shows the overall trend for each tract, averaged across datasets. To the right of each age effect plot, average age effects for superficial (filled circle) and deep (open circle) tract regions are shown for each dataset. Significant differences between age effects of superficial and deep tract regions were assessed using a network enrichment significance test. Stars denote significance levels following FDR correction.

We next examined age effects of mean diffusivity in association tracts and found highly convergent results (**Figure 3**). Similar to callosal tracts, we found that across datasets, most association tracts exhibited the same deep-to-superficial gradient of age effects. However, there were two exceptions: the inferior longitudinal fasciculus and uncinate fasciculus did not exhibit a significant deep-to-superficial patterning as determined by enrichment testing. Nonetheless, 13 out of 15 callosal and bilateral association tracts exhibited significant differences between deep and superficial tract regions in at least two of the three datasets. Furthermore, the magnitude of these age effects were highly correlated across datasets (PNC–HCP-D: *r* = 0.86, *p*_perm_ < 0.0001; PNC–HBN: *r* = 0.72, *p*_perm_ < 0.0001; HCP-D–HBN: *r* = 0.63, *p*_perm_ < 0.0001; **Figure S1**).

**Figure 3.**
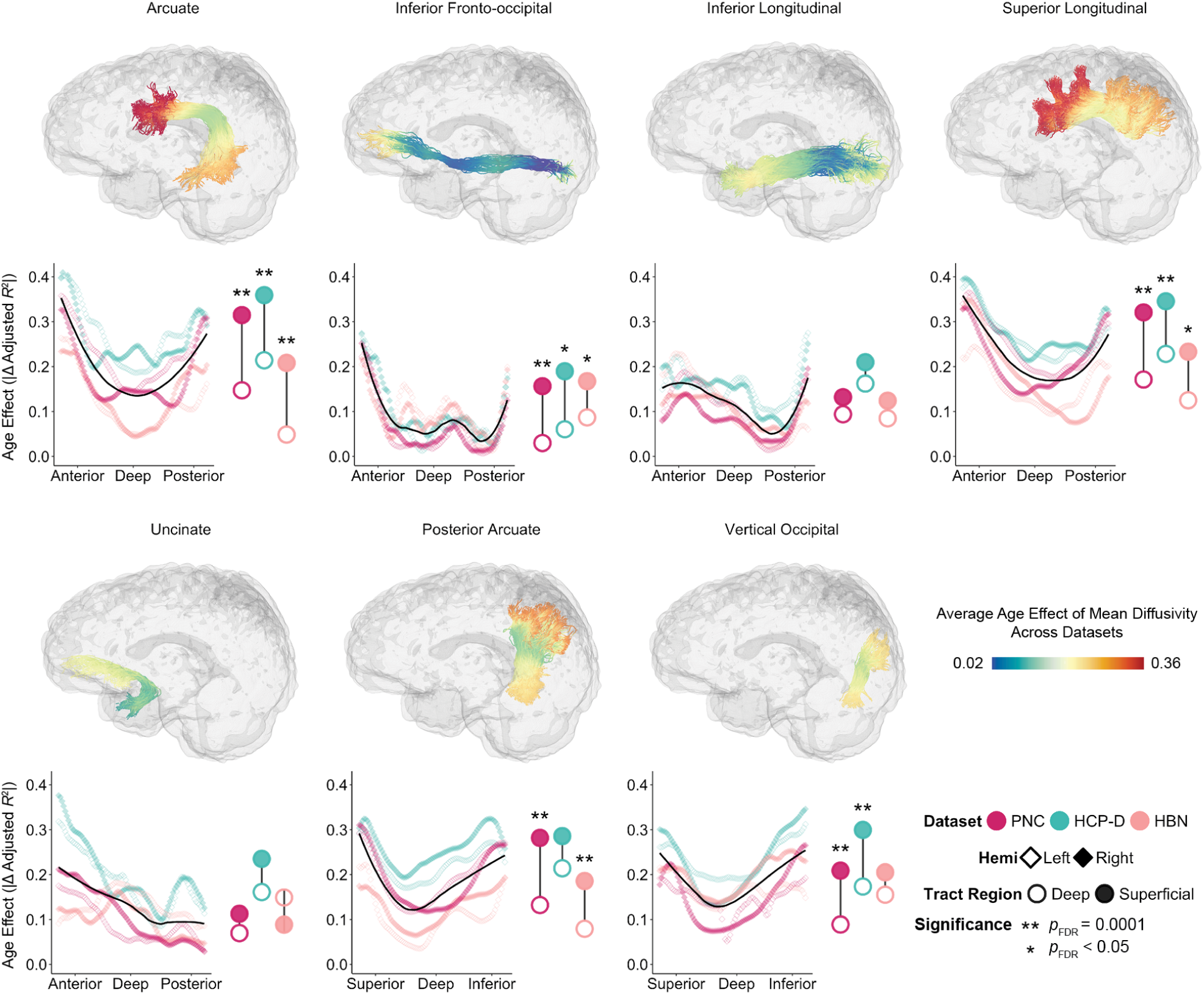
White matter development follows a deep-to-superficial axis in association tracts. Magnitudes of the mean diffusivity age effect along bilateral association tracts vary along a deep-to-superficial axis in PNC, HCP-D, and HBN. Each tract from an exemplar participant is colored by the average magnitude of the mean diffusivity age effect across datasets, depicting the spatial distribution of age effects. Below each glass brain, we display the magnitude of the age effect at 100 equidistant points along the tract’s length. Colors indicate dataset; nodes that do not display significant age effects are shown in gray (*Q*_FDR_ > 0.05). Open and closed diamond shapes represent left and right hemisphere tracts, respectively. The black LOESS-smoothed line shows the trend for each tract averaged across datasets and hemispheres. To the right of each age effect plot, average age effects for bilateral superficial (filled circle) and deep (open circle) tract regions are displayed for each dataset. Statistical comparisons between age effects of superficial and deep tract regions were assessed using a network enrichment significance test, with significance indicated by stars following FDR correction.

To evaluate whether larger age effects in superficial regions of a tract could result from increased variability at the tract ends – providing more variance for age to explain – we compared the coefficient of variation (CV) of mean diffusivity between deep and superficial tract regions (**Table S1**). The CV at superficial tract regions for most tracts appeared to be lower than that in deep tract regions, suggesting that greater magnitudes of superficial age effects were not driven by increased variability in the measure under study. Furthermore, to investigate whether a deep-to-superficial pattern of development generalizes beyond cortico-cortical WM tracts, we additionally evaluated the corticospinal tract (**Figure S2)**. We compared superficial tract regions (adjacent to the motor cortex) to deep tract regions (near the brainstem). As in cortico-cortical tracts, we found that superficial tract regions were significantly enriched for age effects compared to deep WM, suggesting that this pattern does not depend on tract geometry or anatomical orientation. Together, these results describe a highly replicable deep-to-superficial axis of WM development that has previously been obscured by typical methods that average across the length of WM tracts.

### Development of superficial tract regions varies by cortical endpoint similarity

The above results emphasize that superficial tract regions exhibited greater magnitudes of age effects than deep tract regions. However, age effects in superficial tract regions varied both within and across tracts. To investigate this variance, we next evaluated whether variation in age effects in superficial WM was associated with their nearest cortical endpoints. We hypothesized that superficial tract regions would show similar developmental patterns in a tract with similar endpoints, such as homotopic cortices – e.g., corresponding cortical regions in opposite hemispheres. In contrast, we hypothesized that in a tract that linked different cortical endpoints (e.g. heterotopic endpoints), superficial tract regions would display diverging developmental patterns. To test this hypothesis, we developed a workflow that allowed us to create a tract-to-cortex probability map for each tract in each dataset. This workflow overcomes technical obstacles by combining toolkits (e.g., pyAFQ, MRTrix, FSL, Freesurfer, Nilearn, sMRIPrep, and Connectome Workbench), thereby allowing us to relate WM development to each tract’s cortical endpoints. We parcellated the endpoint probability maps using the HCP-MMP atlas.

We first compared the magnitude of age effects for the most superficial nodes in one tract with homotopic endpoints: the motor segment of the corpus callosum (**Figure 4**; i.e., callosum motor). This tract connects homotopic right and left motor cortices. Across all three datasets, we found that superficial regions in this tract exhibited highly similar age effect magnitudes for the WM adjacent to each endpoint (**Figure 4a-c**). Enrichment testing confirmed that the right and left superficial tract regions’ age effects did not significantly differ from each other (**Figure 4d-f)**. We next compared the nonlinear GAM-derived developmental fits of the superficial nodes for each end of the callosum motor. Superficial tract regions adjacent to the right and left motor cortices showed similar patterns of age-related change for all datasets **(Figure 4g-i)**. Right and left motor WM also displayed similar windows of significant developmental change, slowing in developmental change at nearly the same time.

**Figure 4.**
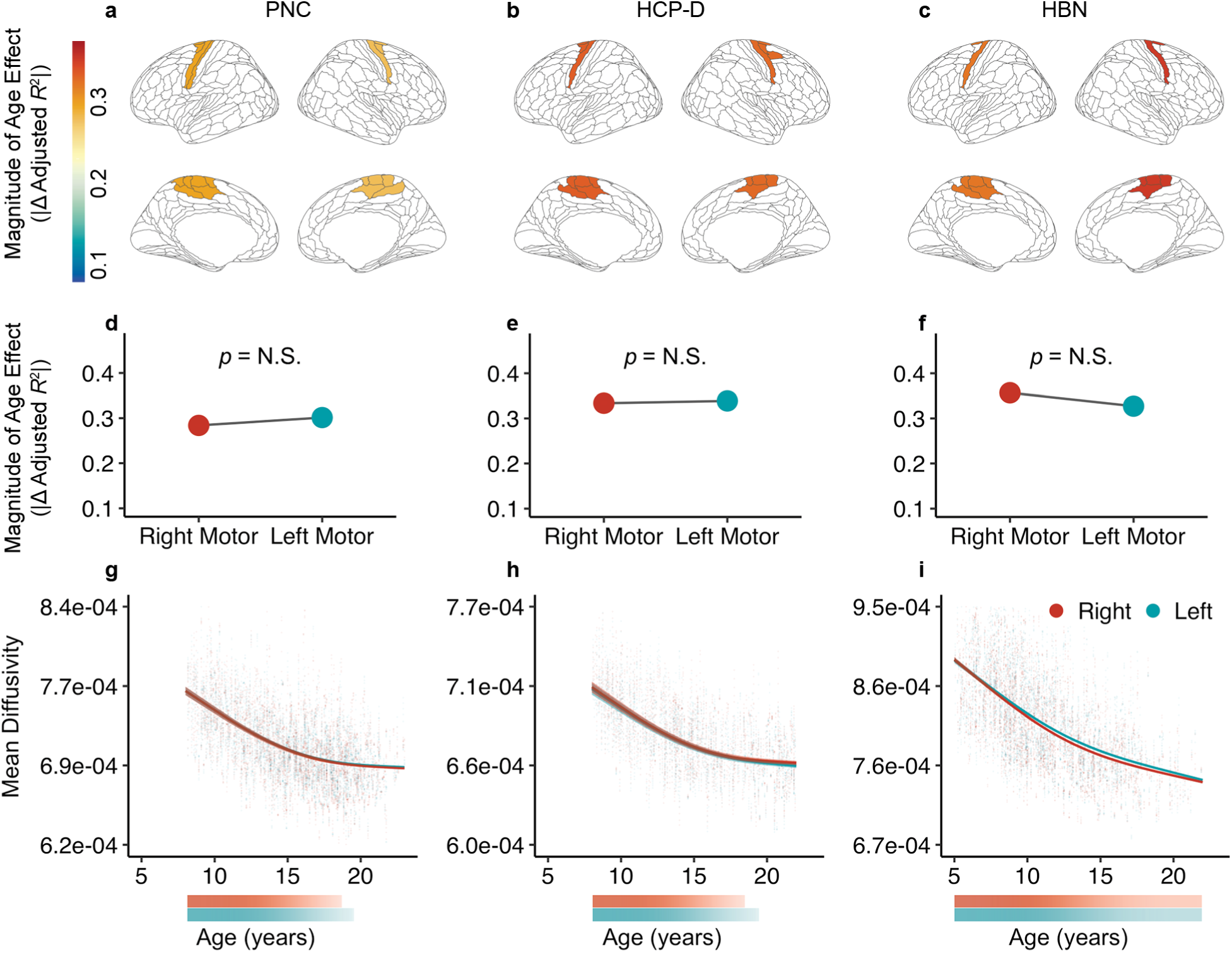
Superficial tract regions in the motor segment of the corpus callosum connecting homotopic cortical endpoints exhibit similar developmental patterns. Developmental measures of superficial tract regions in PNC (first column), HCP-D (second column), and HBN (third column) show similar patterns for each endpoint of the callosum motor. **(a-c)** Magnitudes of the mean diffusivity age effect for the superficial tract regions adjacent to the right and left hemisphere cortical endpoints of the callosum motor are displayed on the cortical surface. **(d-f)** Magnitudes of the age effect at the right (red) and left (blue) motor endpoints do not significantly differ in all three datasets. Statistical comparisons between right and left white matter age effects were assessed using a network enrichment significance test. **(g-h)** Developmental patterns for superficial tract regions are overlaid on participant-level mean diffusivity values. In all three datasets, superficial tract regions adjacent to right and left motor cortices exhibit highly similar patterns. Colored bars indicate windows of significant developmental change in mean diffusivity, with higher transparency corresponding to slower rates of change.

We next investigated developmental patterns at the end of a tract connecting heterotopic endpoints, the inferior fronto-occipital fasciculus (**Figure 5**; IFOF). This tract connects frontal and occipital cortices. Notably, superficial tract regions at these two ends of the IFOF showed dissimilar age effects **(Figure 5a-c)**. Specifically, age effects of frontal WM significantly differed from that of occipital WM in the PNC and HCP-D but not in HBN **(Figure 5d-f)**. In PNC and HCP-D, the magnitude of the age effect was greater in bilateral frontal WM than bilateral occipital WM. When comparing the developmental fits of superficial tract regions in the IFOF, we found that mean diffusivity in occipital WM showed a developmental plateau in mid-adolescence **(Figure 5g-i)**. However, mean diffusivity in frontal WM did not slow in developmental change by the maximum age studied in all datasets, suggesting that it continued to significantly decrease beyond the age window. Interestingly, the cortical regions comprising the frontal endpoint of IFOF sit close to the apex of the cortical hierarchy defined by the sensorimotor-association (S-A) axis. In contrast, regions in the occipital IFOF endpoints have a low average S-A rank. These results suggest that the development of superficial portions of tracts may align with the position of each tract’s endpoint on the cortical hierarchy.

**Figure 5.**
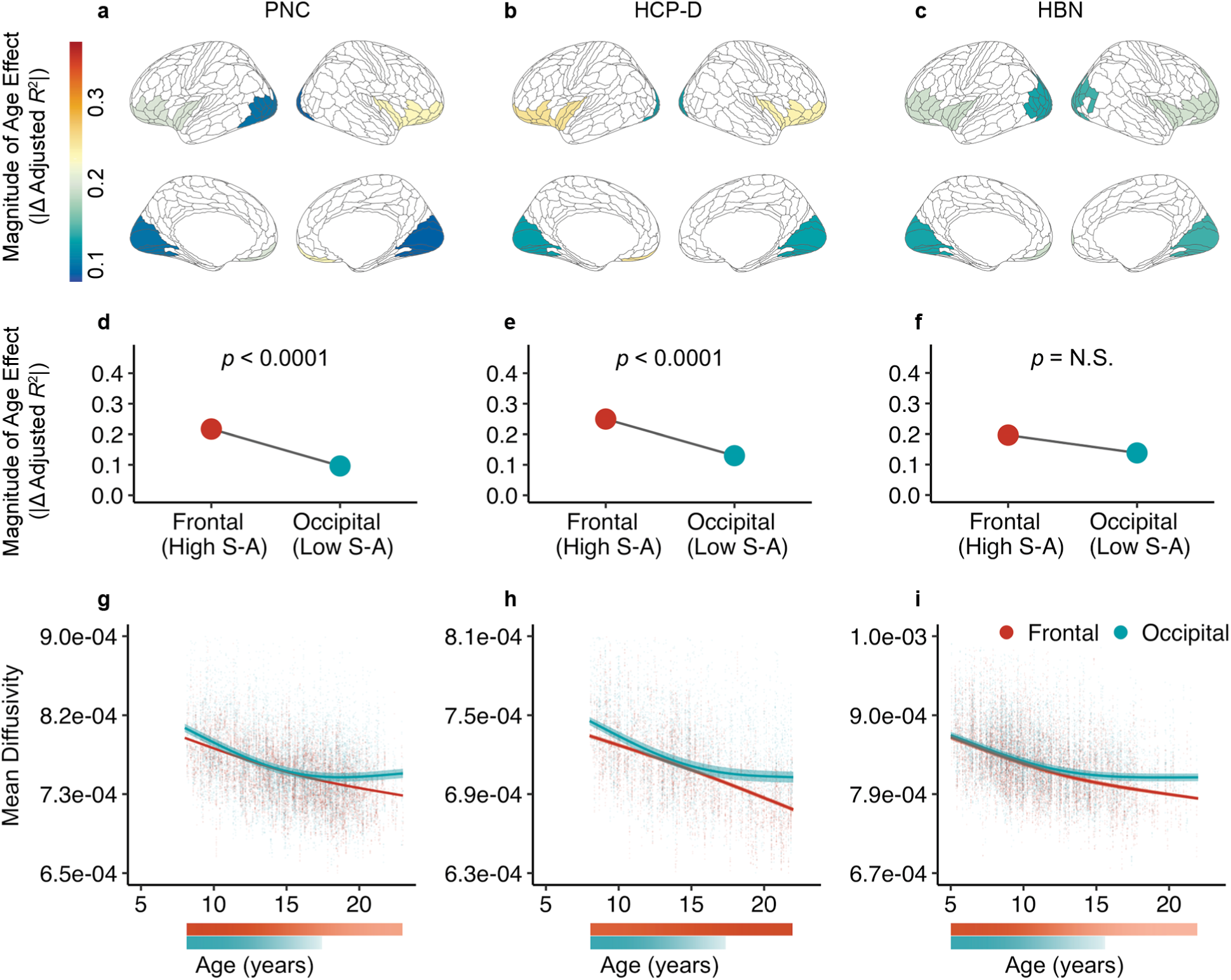
Superficial tract regions in inferior fronto-occipital fasciculus connecting heterotopic cortical endpoints exhibit distinct developmental patterns. Superficial tract regions in the inferior fronto-occipital fasciculus (IFOF) exhibit distinct developmental patterns between the frontal and occipital endpoints, which respectively have high and low sensorimotor-association (S-A) ranks on average. **(a-c)** The magnitudes of the age effect for frontal and occipital white matter of IFOF are shown on the cortical surface. **(d-f)** The age effect, averaged across hemispheres, is significantly larger in frontal (red) compared to occipital (blue) white matter in PNC and HCP-D. Statistical comparisons between frontal and occipital white matter age effects were assessed using a network enrichment significance test. **(g-h)** Developmental patterns for frontal and occipital white matter are overlaid on participant-level mean diffusivity values. Colored bars depict windows of significant developmental change, with higher transparency indicating slower rates of change. In all datasets, occipital white matter matures in mid-adolescence whereas frontal white matter continues to significantly decrease in mean diffusivity beyond the studied age window.

### Development of superficial tract regions aligns with the sensorimotor-association axis

Contrary to the notion that tracts develop synchronously along their entire length, the above findings reveal that within a single tract, superficial tract regions adjacent to heterotopic cortical endpoints may diverge in maturational timing. In the IFOF, such divergence appeared to be consistent with the cortical hierarchy. We hypothesized that the relative positions of each tract’s cortical endpoints along the cortical hierarchy would explain the observed convergence or divergence in their developmental patterns. Thus, we expanded this analysis to all tracts and interrogated whether the age of maturation for superficial tract regions aligned with each endpoint’s position on the S-A axis. Given that cortical maturation has been shown to develop along the S-A axis, we specifically hypothesized that microstructure in superficial tract regions would exhibit analogous developmental patterns, with unimodal, lower-order regions maturing earlier and transmodal, higher-order regions maturing later.

To evaluate this hypothesis, we computed the age of maturation as the earliest age at which the rate of developmental change was no longer statistically different than zero. We then averaged the ages of maturation for the most superficial nodes for each tract. We computed the correlation between the ages of maturation at each endpoint with the mean S-A axis rank of that endpoint’s constituent cortical regions. We found that in two of the three datasets, ages of maturation for each endpoint were largely explained by each endpoint’s mean S-A axis rank (**Figure S3a-d**; PNC: *r* = 0.81, *p*_spin_ < 0.0001; HCP-D: *r* = 0.85, *p*_spin_ < 0.0001; HBN: *r* = 0.04, *p*_spin_ = N.S.). When averaged across datasets, a significant effect remained (*r* = 0.67, *p*_spin_ < 0.0001). Furthermore, comparing the age of maturation and the magnitude of the mean diffusivity age effect revealed a moderate correlation, suggesting that while these measures capture overlapping aspects of development, they also provide distinct developmental information (**Figure S4a-c;** PNC: *r* = 0.59, *p*_perm_ < 0.0001; HCP-D: *r* = 0.43, *p*_perm_ < 0.0001; HBN: *r* = 0.46, *p*_perm_ < 0.0001).

It should be noted that the cortical endpoints of many tracts did not mature by the maximum studied age in each dataset, resulting in a ceiling effect as these endpoints were assigned the maximum age studied in each dataset. To evaluate the potential influence of ceiling effects, we included only the endpoints that reached maturation within the age window of our samples in a sensitivity analysis. This analysis generated consistent results: the age of maturation was associated with S-A rank in PNC and HCP-D (**Figure S3e-h**; PNC: *r* = 0.77, *p*_spin_ < 0.0001; HCP-D: *r* = 0.5, *p*_spin_ < 0.05; HBN: *r* = 0.28, *p*_spin_ < 0.05; averaged across datasets: *r* = 0.7, *p*_spin_ < 0.0001). Additionally, to determine whether still-developing endpoints had higher S-A ranks than endpoints that were no longer significantly changing in mean diffusivity, we conducted a two-sample, one-tailed spin-based *t*-test. In the PNC, late-developing endpoints had significantly higher S-A ranks than endpoints that matured in the sample age range (*p*_spin_ < 0.0001). This finding was replicated in HCP-D (*p*_spin_ < 0.0001) but not in HBN.

To further explore hierarchical development between superficial regions within each tract, we examined whether tracts that spanned the cortical hierarchy exhibited greater differences in their age of maturation compared to tracts that connected similar regions in the cortical hierarchy. To do this, we compared differences in age of maturation (ΔAge) of tracts with large differences in S-A axis rank (ΔS-A rank) between a given tract’s endpoint to those with a small ΔS-A rank. Callosal tracts connecting similar cortical regions, such as callosum motor **(Figure 6a)** and several association tracts, including arcuate fasciculus and superior longitudinal fasciculus, displayed small differences in mean ΔS-A rank (generally < 50).

**Figure 6.**
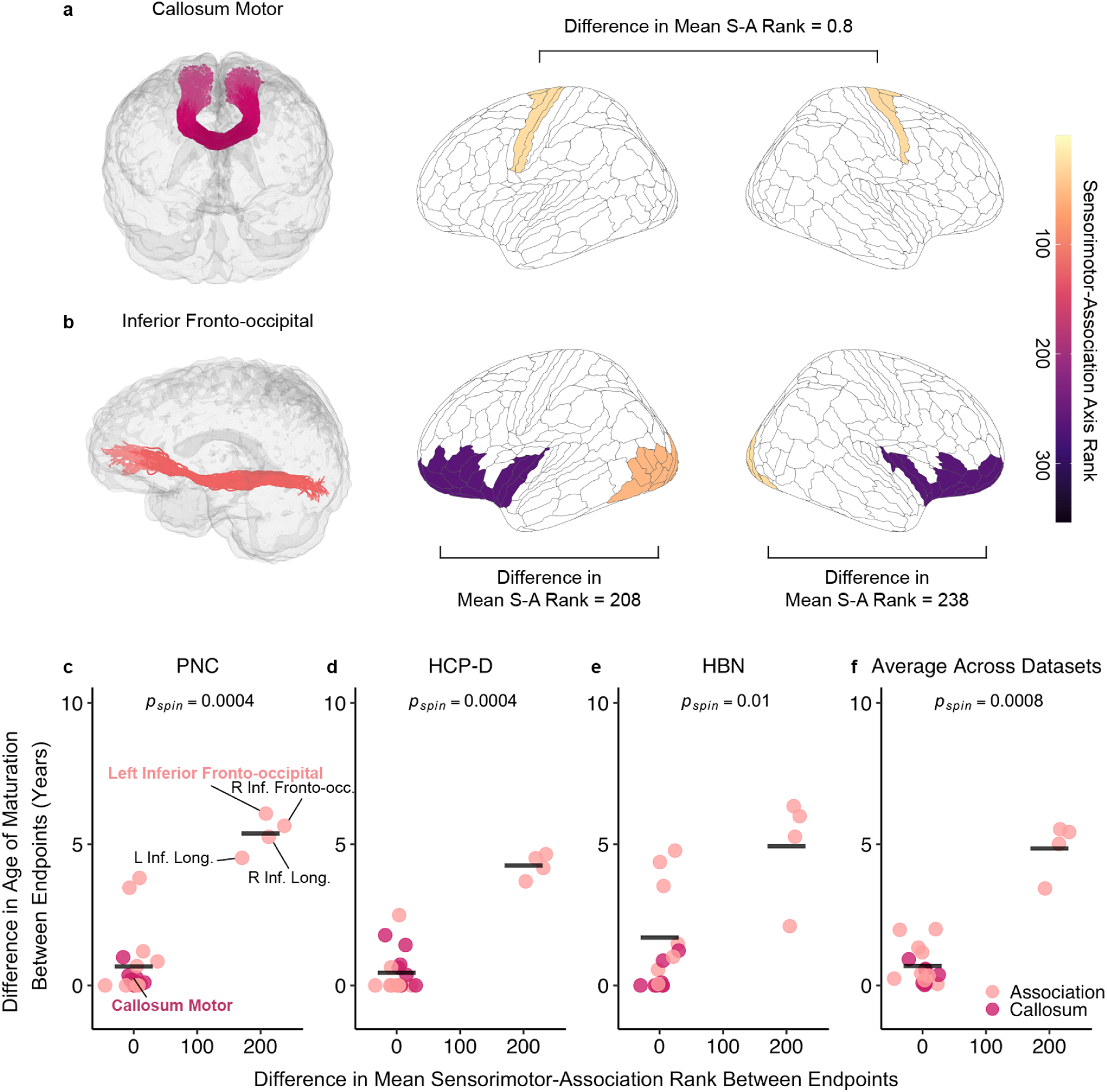
Cortical endpoints at opposite ends of the cortical hierarchy have discrepant ages of maturation. **(a)** Callosum motor terminates on homotopic motor regions, resulting in a small mean S-A rank difference of 0.8. **(b)** The inferior fronto-occipital fasciculus is a long-range association tract that connects frontal regions with high sensorimotor-association (S-A) axis ranks to occipital regions with low S-A ranks, resulting in a large mean S-A rank difference (>200) between its endpoints in each hemisphere. **(c-e)** The relationship between the difference in age of maturation (ΔAge) and the difference in mean S-A axis rank (ΔS-A rank) between endpoints is displayed for **(c)** the PNC, **(d)** HCP-D, **(e)** HBN, and **(f)** across datasets. Each data point represents a unique white matter tract, colored by tract type (association tracts in light pink, callosal tracts in dark pink). Tracts connecting regions with small ΔS-A rank—including all callosal tracts and several association tracts such as the arcuate—exhibit small ΔAge. In contrast, bilateral inferior fronto-occipital and inferior longitudinal fasciculi, which connect regions with large ΔS-A rank, exhibit large ΔAge between their endpoints. In all datasets, spin-based permutation tests confirmed that tracts with large differences in S-A rank have significantly greater ΔAge between their endpoints compared to tracts with small differences. Horizontal lines indicate the mean ΔAge for the two groups.

Long-ranging tracts connecting high- to low-order regions included the IFOF **(Figure 6b)** and inferior longitudinal fasciculus and displayed a large ΔS-A rank (∼200). This framework broadly revealed two groups of tracts: tracts connecting hierarchically similar regions with similar maturational ages, and tracts spanning the hierarchy with discrepant ages of maturation **(Figure 6c-f)**. For example, all callosal tracts and several association tracts had highly similar ages of maturation (i.e., small ΔAge). In contrast, bilateral IFOF and inferior longitudinal fasciculus exhibited a large discrepancy in ages of maturation between their endpoints (i.e., large ΔAge). A one-tailed, spin-based *t*-test revealed that tracts with larger ΔS-A rank had greater ΔAge between endpoints across all datasets (**Figure 6c-f**; PNC: *p*_spin_ = 0.0004; HCP-D: *p*_spin_ = 0.0004; HBN: *p*_spin_ = 0.01; averaged across datasets: *p*_spin_ = 0.0008). These findings underscore that the superficial regions of tracts that connect similar regions along the cortical hierarchy mature at a similar age. In contrast, superficial regions of tracts that traverse the cortical hierarchy have discordant ages of maturation.

To summarize broad developmental patterns of superficial tract regions in the context of the cortical hierarchy, we extended our within-tract analyses to study developmental variation across tracts. For each dataset, we aggregated the ages of maturation for superficial tract regions across all tracts and assigned the average age of maturation to the nearest cortical endpoint. This process yielded an average age of maturation map across all superficial tract regions, where each HCP-MMP region that had a tract termination was assigned an average age of maturation.

Endpoints that matured beyond the maximum age studied in each dataset were initially not included. We found that the age of maturation for each HCP-MMP region with a matured tract termination significantly correlated with the S-A axis in each dataset and also across datasets (**Figure 7a-d**; PNC: *r* = 0.76, *p*_spin_ < 0.0001; HCP-D: *r* = 0.4, *p*_spin_ = 0.0025; HBN: *r* = 0.56, *p*_spin_ < 0.0001; averaged across datasets: *r* = 0.58, *p*_spin_ < 0.0001). As prior, to determine whether late developing endpoints had higher S-A ranks than endpoints exhibiting evidence of maturation, we conducted a two-sample, one-tailed *t*-test with a spin test. In the PNC, still-developing endpoints had significantly higher S-A ranks than mature endpoints (*p*_spin_ < 0.0001). This finding was replicated in HCP-D (*p*_spin_ < 0.0001) but not in HBN. Additional analyses that included all endpoints (regardless of when they matured) found that age of maturation was significantly associated with S-A axis rank in PNC and HCP-D (**Figure S5**; PNC: *r* = 0.72, *p*_spin_ < 0.0001; HCP-D: *r* = 0.77, *p*_spin_ < 0.0001; HBN: *r* = 0.23, *p*_spin_ = 0.006; averaged across datasets: *r* = 0.54, *p*_spin_ < 0.0001). Taken together, these results demonstrate that superficial tract regions develop hierarchically along the S-A axis.

**Figure 7.**
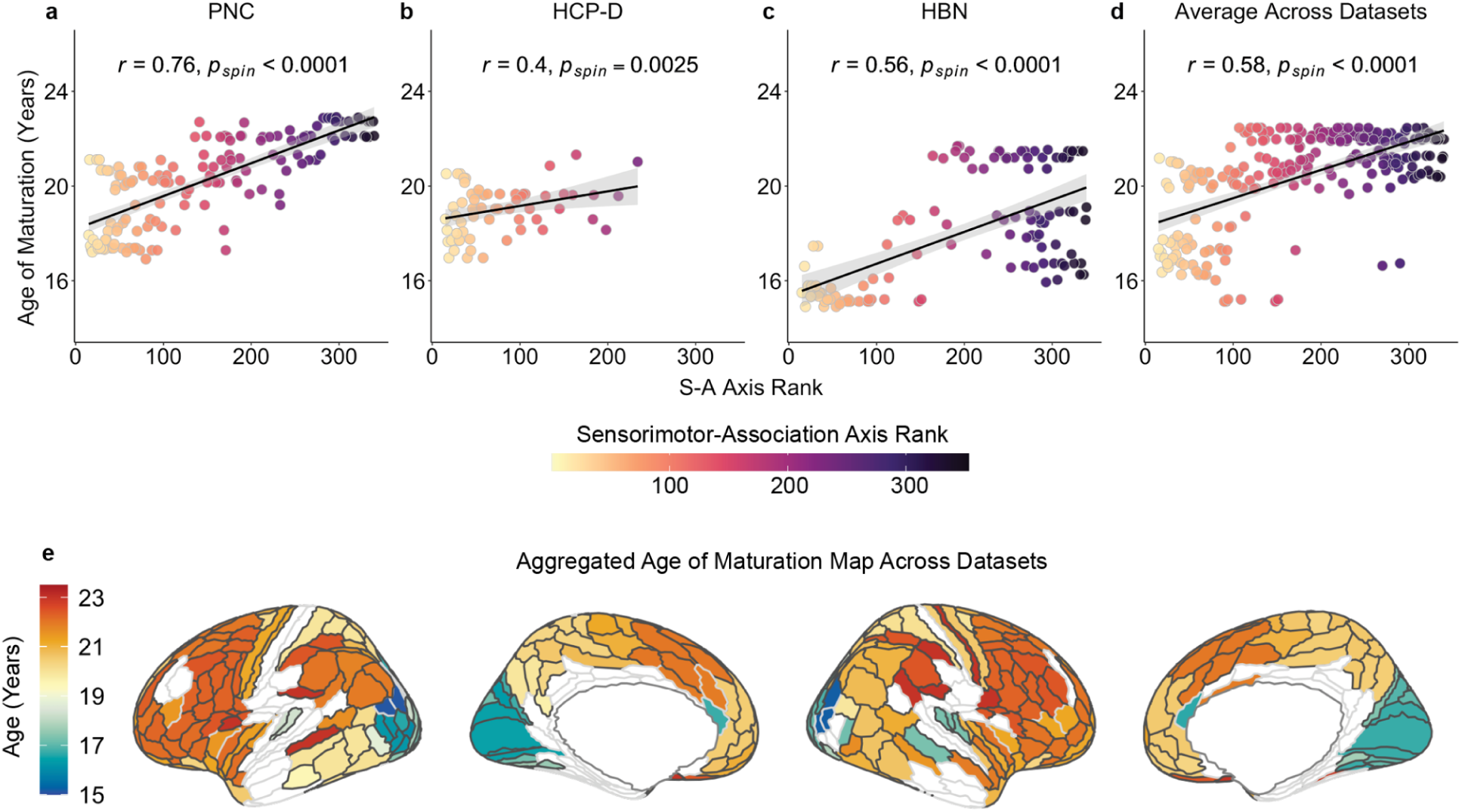
Superficial tract regions develop hierarchically along the sensorimotor-association axis. Parcellated cortical maps were created by averaging ages of maturation across superficial tract regions and assigning the average value to each HCP-MMP region that had a tract termination for each dataset. **(a-c)** Age of maturation is associated with S-A axis rank in each dataset; regions with older ages of maturation rank higher on the S-A axis in all three datasets: **(a)** PNC (*r* = 0.76, *p*_spin_ < 0.0001), **(b)** HCP-D (*r* = 0.4, *p*_spin_ = 0.0025), and **(c)** HBN (*r* = 0.56, *p*_spin_ < 0.0001), as well as **(d)** across all datasets (*r* = 0.58, *p*_spin_ < 0.0001). Statistical significance of correlations was assessed using region-based spin tests. This analysis excluded superficial tract regions that matured later than the maximum age studied in each dataset; see **Figure S5** for similar results that include these regions. **(e)** Dataset-specific maps were then averaged to produce a cross-dataset map. White represents HCP-MMP regions without a tract termination.

## DISCUSSION

We identified two major axes of human white matter (WM) development. First, by examining developmental variation along tracts, we delineated a robust and highly replicable deep-to-superficial axis of WM development, characterized by greater age-related change in superficial compared to deep tract regions. This pattern has not been systematically quantified along major WM tracts with *in vivo* human neuroimaging, as prior studies have focused on discrete WM sites^14,15^ or whole-tract averages^2,8,9,32^. In addition to identifying variation in WM development along the full length of tracts, we studied maturational variability between each tract’s cortical endpoints. Specifically, to study how tracts develop in relation to the cortical regions they connect, we created a methodological workflow to map tracts to their cortical endpoints. This advance allowed us to demonstrate that the maturation of superficial tract regions varies by cortical endpoint similarity. Superficial regions in tracts connecting similar cortical endpoints followed similar developmental patterns whereas the development of endpoints in tracts that spanned the hierarchy diverged. Specifically, we showed that the maturation of superficial tract regions was heterochronous and varied along the cortical hierarchy as defined by the sensorimotor-association (S-A) axis^3^. This work reveals that the spatial and temporal patterning of WM development occurs along two axes: a deep-to-superficial axis and the S-A axis. As discussed below, these co-evolving developmental programs pair early maturation in deep tract regions that may be crucial for signaling fidelity with hierarchical maturation of superficial tract regions that may allow for protracted activity-dependent refinements in youth.

Across nearly all tracts and datasets, we found larger developmental effects in superficial as compared to deep tract regions. This discrepancy suggests that deeper WM regions may be relatively further along in their maturational course compared to superficial tract regions within the studied age range. Such spatial variation in WM maturation has been previously reported in both animal and human studies. In rodents, myelin basic protein first stains deeper WM structures near the brainstem during the neonatal stage before extending outward and rostrally toward the cortex during the juvenile period^33^. Post-mortem studies in infants have described earlier myelination of deep WM structures (e.g., the posterior and anterior limbs of the internal capsule and body, splenium, and rostrum of the corpus callosum) compared to WM regions near the cortex such as the frontal and temporal poles^14,15^. Similarly, small-scale human neuroimaging studies in infancy and early childhood have provided results that align with studies in rodents, with diffusion signal in deep WM structures showing larger developmental change earlier than WM adjacent to cortex^33–36^. Notably, a recent along-tract neuroimaging study of infants found faster development of deep tract regions^37^. These findings from animal systems and early development in humans dovetail with our findings in childhood and adolescence, where deep WM showed relatively little change. Instead, associations with age were of greater magnitude in superficial tract regions – a pattern that was consistently observed across diverse tracts. Together, our findings and prior literature suggest that WM tract development is heterochronous: deeper WM tract regions undergo more developmental change earlier, while superficial tract regions continue to develop into childhood and adolescence.

Why might there be a prominent deep-to-superficial axis of WM development? We speculate that early development of deep tract regions may reflect earlier myelination that is essential for enabling fidelity in neural transmission. The first wave of myelination in the brain – marked by a rapid onset of dramatic myelin changes^38^ – begins around birth and continues through the first years of life. Myelin formation is thought to be driven by two processes: the intrinsic (activity-independent) and adaptive (activity-dependent) pathways^38,39^. In this initial wave of myelination, activity-independent processes may predominate^38,40,41^, with myelination driven by deep WM oligodendrocytes that have unique transcriptional programs^42^. For example, oligodendrocytes are several-fold denser in WM than in gray matter^43^, where their expansion occurs later in development^39,44^. Although research remains sparse on regional differences in oligodendrocyte transcriptional programs within WM, we speculate that dense populations of oligodendrocytes in deep WM may be genetically programmed for rapid early myelination.

Activity-dependent mechanisms may also contribute to preferential myelination in deep WM during early development. Developmental studies in rodents and zebrafish have shown that neuronal activity leads to proliferation of oligodendrocyte progenitor cells (OPCs), differentiation of OPCs into oligodendrocytes, and preferential ensheathment of active axons^17,45^. While activity-dependent myelination is largely specific to active axons, increases in myelination also occur on nearby axons through a “bystander effect”^45^, possibly mediated by axonal signaling factors that locally regulate myelin^17^. Deep tract regions are characterized by a smaller tract radius with more densely packed axons than superficial tract regions that fan out to the cortex^46^. Because of this compact arrangement, axons in deep tract regions may be particularly susceptible to the bystander effect, leading to disproportionate increases in myelination during infancy and early childhood. We hypothesize that the greater developmental changes observed in superficial tract regions compared to deep tract regions may be due to the latter having undergone much of their myelination prior to the age ranges we studied here – as early as infancy^37^ – bringing them closer to a developmental plateau by childhood and adolescence. We note that while mean diffusivity is not a direct marker of myelin and reflects multiple microstructural properties, our developmental framework aligns with prior literature highlighting the role of myelination in WM development. Furthermore, we replicated developmental findings using multi-shell measures from NODDI and MAP-MRI, which have been validated by histology^47^ and magnetization transfer imaging^48^ to be sensitive to myelin in white matter. Early preferential myelination of deep tract regions may serve to insulate axons from their neighbors and reduce unwanted ephaptic coupling (electrical crosstalk between adjacent axons)^49^, establishing the basis for successful transmission of action potentials between distant cortical regions. This first wave of myelination may set the stage for the subsequent developmental refinement at superficial tract regions in childhood through early adulthood that we observe here.

This epoch of childhood, adolescence, and young adulthood coincides with increasingly complex cognitive, social, and emotional environmental inputs^50^, a protracted period of spatially heterogeneous refinements in intrinsic cortical activity^5^, and a second wave of rapid myelin changes^38^. These experiences, and associated neural remodeling, may contribute to the pronounced changes in superficial tract regions seen in our study. In juvenile rodents, after deep WM has heavily myelinated, neuronal activity preferentially drives activity-dependent changes in OPCs in the deep layers of cortex and the adjacent white matter (i.e. superficial tract regions) compared to deep WM^51^. Consistent with this patterning, developmental change in superficial tract regions in childhood and adolescence may reflect continued remodeling in WM regions proximate to neuronal activity that may result from factors including novel life experiences^50^ and intrinsic cortical activity^5^. Marked changes in superficial tract regions may be driven in part by glutamate release and growth factor signaling in the cortex, which facilitate activity-dependent myelination^38^. Similarly, nearby neuronal cell bodies release OPC mitogens, promoting local OPC proliferation and subsequent myelination^51^. Furthermore, deep layers in the developing frontal cortex have been shown to myelinate earlier than superficial layers^52^; our results extend this deep-to-superficial maturational sequence to long-range WM tracts. Together, our findings and the existing literature suggest that the overall myelination of the brain may exhibit an “inside-out” developmental pattern that begins in deep WM, extends into superficial tract regions and deep cortical layers, and finally reaches the cortex’s outermost layers. Ultimately, our results are consistent with a model in which experience-driven cortical changes and ongoing cortical refinement dynamically interact with the remodeling of superficial tract regions near cortex in childhood and adolescence^38^.

In addition to this deep-to-superficial axis of development, we also found that age-related changes in superficial tract regions did not occur uniformly and instead varied between cortical endpoints within a single tract. For example, the inferior fronto-occipital fasciculus spans frontal to occipital regions; we observed an older age of maturation in frontal WM and earlier maturation in occipital WM. Furthermore, our findings not only showed differential developmental timing along individual tracts based on their cortical endpoints, but also that the development of superficial tract regions aligned with the cortical hierarchy as defined by the S-A axis^3^. These results provide empirical evidence for a hypothesis first proposed more than a century ago by Flechsig^53,54^ and restated by Yakovlev and Lecours^14^ that myelination proceeds along a hierarchy of increasingly complex cortical functions^15^. The development of cortical myelination has been shown to align with the S-A cortical hierarchy^6^, with intracortical myelination exhibiting protracted developmental changes and later maturation in higher-order association regions^6,7,55^. Building on this existing literature on intracortical myelin, we show that superficial tract regions of long-range cortico-cortical tracts also develop heterochronously, progressing from sensorimotor regions to higher-order association regions. Hierarchical refinements in myelin may help fine-tune the precise timing of action potentials^13,56^, which may facilitate synchronization of long-range information transmission^57^ and optimize cortical oscillations important for complex cognitive functions^57,58^. Together, our results challenge the notion that a tract matures uniformly along its length, revealing that along-tract development may be governed in part by the position of a tract’s endpoints on the cortical hierarchy.

This study has several important limitations. First, we examined WM microstructure using advanced methods in diffusion modeling and tractography, which do not directly represent axons. Specifically, our primary measure was mean diffusivity, which can be influenced by the presence of astrocytes, microglia, and oligodendrocytes as well as myelin^23^. Second, superficial tract regions that fan out to the cortex may introduce partial volume effects, though mean diffusivity is less susceptible to these effects and the use of multi-shell acquisition and anatomically-constrained tractography helps to further mitigate them. Third, we specifically examined the major association and commissural pathways, including one projection pathway (the corticospinal tract) as a sensitivity analysis. Thus, we did not examine short-range association U-shaped fibers. Fourth, our age window precludes us from examining development in infancy and early childhood, when dramatic WM changes occur. Similarly, the oldest subjects we include were age 23, leading to ceiling effects in computing the age of maturation in superficial tract regions. Of note, many regions that were impacted by the age ceiling effect ranked relatively high on the S-A axis. Lastly, WM has been classically described to develop from posterior to anterior regions. However, the posterior-anterior axis is difficult to disambiguate from the hierarchical S-A axis as the two axes are highly correlated, especially without full coverage of the cortex.

In this study, we replicably identified two distinct axes of development along major WM tracts in the human brain. First, we demonstrated that WM develops along a deep-to-superficial axis. Second, we demonstrated that development of superficial tract regions progresses hierarchically across tracts along the S-A axis. By examining development along the continuous length of tracts rather than relying on discrete WM regions or tract averages, our results provide a more complete account of WM development. This work argues against the implicit assumption of synchronous maturation along tracts – an assumption reinforced by methodological constraints of conventional tract-average analyses – and underscores the need to consider variable development along tracts when characterizing WM maturation. Understanding the biological mechanisms driving the heterochronous, hierarchical refinement of WM requires complementary work in animal models using causal experimental designs, where molecular processes influencing myelination can be directly examined. In humans, future work using MRI techniques that are sensitive to myelin, such as myelin water imaging, magnetization transfer imaging, and T1 relaxometry^59^ in longitudinal samples would allow investigation into the developmental progression of myelin within individuals. Moving forward, the recognition that WM tracts are not uniform conduits between cortical regions but exhibit developmental variability along their length will deepen our understanding of how WM and cortical development are intertwined. Ultimately, this work will help us better characterize human brain organization and variation associated with neuropsychiatric illness across the lifespan.

### **STAR** ★ **METHODS**

### KEY RESOURCES TABLE

| REAGENT or RESOURCE | SOURCE | IDENTIFIER |
| --- | --- | --- |
| **Deposited data** |  |  |
| The Philadelphia Neurodevelopmental Cohort |  | ncbi.nlm.nih.gov/projects/gap/cgi-bin/study.cgi?study_id=phs000607.v3.p2 |
| The Lifespan Human Connectome Project Development (HCP-D) | Somerville et al. | humanconnectome.org/study/hcp-lifespan-development |
| Healthy Brain Network | Alexander et al. | fcon_1000.projects.nitrc.org/indi/cmi_healthy_brain_network |
| **Software and algorithms** |  |  |
| pyAFQ | Kruper et al. | tractometry.org/pyAFQ |
| QSIprep | Cieslak et al. | qsiprep.readthedocs.io/en/0.22.0 |
| Freesurfer | Dale et al.<br>Fischl et al. | surfer.nmr.mgh.harvard.edu/ |
| R | The R Project | r-project.org |
| Python | Python Software Foundation | [Python.org](https://python.org) |
| BABS | Zhao et al. | github.com/PennLINC/babs/releases/tag/0.0.8 |
| Tract-to-cortex mapping | This paper | github.com/PennLINC/luo_wm_dev/tree/main |

#### STUDY PARTICIPANT DETAILS

Analyses were conducted in three large-scale datasets. The Philadelphia Neurodevelopmental Cohort (PNC; *n* = 1,098) was the discovery dataset, while the Human Connectome Project: Development (HCP-D; *n* = 567) and Healthy Brain Network (HBN; *n* = 1,051) served as replication datasets. Study procedures in each dataset were approved by the following Institutional Review Boards: the University of Pennsylvania and Children’s Hospital of Philadelphia Institutional Review Boards for the PNC, a central Institutional Review Board at Washington University in St. Louis for HCP-D, and the Chesapeake Institutional Review Board (now Advarra Inc.) for HBN. For all datasets, written informed consent was obtained for participants over 18 years of age. Informed consent was provided by legal guardians and informed assent was obtained from participants under 18 years of age. Demographic information for PNC, HCP-D, and HBN are reported in **Table 1**.

**Table 1.** Demographic characteristics for each dataset. Participants self-reported race and sex; intersex was not assessed. The racial category “Other/Mixed” includes individuals who identified with more than one race and those identifying as American Indian or Alaska Native, Hispanic or Latino, or Native Hawaiian or Other Pacific Islander.

| Dataset | N | Female (%) | Age Range (Years; Mean± SD) | Race (self-reported) |  |  |  |  |
| --- | --- | --- | --- | --- | --- | --- | --- | --- |
|  |  |  |  | Asian | Black | Other/Mixed | White | Unknown |
| PNC | 1098 | 579 (52.7%) | 8-23 (15.3±3.5) | 10 (0.9%) | 464 (42.3%) | 115 (10.47%) | 509 (46.36%) | 0 (0%) |
| HCP-D | 567 | 303 (53.4%) | 8-22 (14.8±3.9) | 44 (7.76%) | 59 (10.41%) | 82 (14.46%) | 368 (64.9%) | 14 (2.47%) |
| HBN | 1051 | 386 (36.7%) | 5-22 (11.3±3.6) | 30 (2.9%) | 132 (12.6%) | 282 (26.8%) | 488 (46.4%) | 119 (11.3%) |

The PNC^60^ is a community sample of children and adolescents from the greater Philadelphia area recruited for studying typical and atypical brain development. Minimal initial exclusion criteria were applied to the PNC and included medical conditions that could impact brain function^61^. Data from 1,098 participants ages 8-23 years from the PNC were included in the current study after additional exclusion criteria were applied; see “Sample Construction” for details.

HCP-D^62^ is a sample of typically developing children and adolescents. To reflect the demographics of youth in the U.S., participants were recruited across four academic sites: University of Minnesota, Harvard University, Washington University in St. Louis, and University of California-Los Angeles. Information about initial inclusion and exclusion criteria is described previously, with participants excluded for medical conditions that could impact brain function^62^. After applying additional exclusion criteria to the Lifespan 2.0 release, we included demographic and neuroimaging data from 567 participants ages 8-22 years in the present study.

Lastly, HBN^63^ is a self-referred sample of children and adolescents residing in the New York City area who participated in the study due to concerns about neuropsychiatric symptoms. Data collection took place at four sites: Staten Island Flagship Research Center, Rutgers University Brain Imaging Center, CitiGroup Cornell Brain Imaging Center, and CUNY Advanced Science Research Center. To capture phenotypic heterogeneity in youth psychopathology, HBN’s study exclusion was minimal.^63^ Data from 1,051 participants ages 5-22 from data releases 1-9 in HBN were included in the current study after additional exclusion criteria were applied.

### METHOD DETAILS

#### Sample construction

The following exclusion and quality assurance criteria were applied successively in the order described below and are summarized in **Figure S6**.

##### Variant acquisition exclusion

For the present study, participants who had 3T MRI data with T1w images, field maps, and non-variant acquisitions^64^ of diffusion MRI scans with identical parameters within datasets were considered for inclusion. The following numbers of individuals had all required neuroimaging data: *n* = 1,368 in PNC, *n* = 640 in HCP-D, and *n* = 1,755 in HBN.

##### Medical history exclusion

While all datasets applied an initial medical exclusion (described above), health history exclusion criteria for the current study included presence of additional medical conditions affecting brain function or gross neurological abnormalities. In the PNC, *n* = 118 were excluded from the initial sample of *n* = 1, 368. In HCP-D, *n* = 7 participants were excluded from the initial sample of *n* = 640. No additional health history exclusion was applied to HBN.

##### Imaging protocol quality assurance

T1-weighted images that did not survive manual quality assurance were excluded in HBN and PNC. Participants were further excluded if their raw diffusion scans had missing gradient directions. For the PNC, three highly trained raters visually assessed images and provided manual ratings based on artifacts. *N* = 25 participants were excluded for T1w quality and *n* = 10 participants were excluded for missing gradient directions in the PNC. No additional T1w exclusion was applied to HCP-D, as initial T1w quality assurance was completed by the team that collected the data, but *n* = 8 participants were excluded for missing gradient directions in HCP-D. The Swipes for Science web application^65^ was used to perform manual quality control in HBN.^66^ *N* = 72 participants were excluded for T1w quality in HBN. No participants in HBN were missing gradient directions.

##### dMRI quality assurance

We utilized two measures for dMRI image quality assurance: in-scanner head motion and neighborhood correlation. We excluded participants if their diffusion scans exhibited high in-scanner head motion, as defined as mean framewise displacement > 1 mm. Furthermore, individuals with low neighborhood correlation were excluded. Neighborhood correlation measures the average pairwise spatial correlation between diffusion volumes that sample similar points in *q*-space^67^. Thus, lower values indicate worse data quality. Because neighborhood correlation values vary by diffusion scan acquisition parameters and noise level, the following dataset-specific exclusion thresholds were applied as in previous work: 0.9 in PNC, 0.6 in HCPD, and 0.7 in HBN^68^. In PNC, *n* = 70 participants did not meet the above criteria and were excluded. For HCP-D, *n* = 39 participants were excluded. In HBN, *n* = 373 individuals were excluded.

Because HBN is a higher noise and head motion dataset, we applied additional diffusion quality exclusion provided by HBN Preprocessed Open Diffusion Derivatives (HBN-POD2) quality control criteria^69^. This criteria excludes participants based on expert raters and a convolutional neural network model trained on imaging data and automated quality control metrics. An additional *n* = 208 participants failed HBN-POD2 quality control and were excluded. Spatial variability in data quality, such as greater susceptibility to motion-related artifacts or signal dropout in anterior compared to posterior regions, could potentially confound observed developmental patterns along tracts.

##### Age exclusion

For each dataset, participants ages 5-23 years were included in our study. In the PNC, no additional participants were excluded since all participants were within the age window studied. Sparse sampling of ages < 8 years in this dataset (*n* = 14) may negatively impact accurate developmental modeling of HCP-D^6,68^. Thus, a young age exclusion of participants < 8 years old was additionally applied in HCP-D. In HBN, *n* = 1 participant was excluded for missing age data. A young age exclusion was not applied to HBN, as this dataset included a large sample (*n* > 200) of participants under age 8.

##### Failed reconstruction exclusion

Participants who failed FreeSurfer surface reconstruction or diffusion reconstruction were excluded, which typically results from a low-quality image. In PNC, *n* = 47 participants were excluded. In HCP-D, *n* = 5 participants were excluded. In HBN, *n* = 50 participants were excluded from analyses.

#### MRI data acquisition

PNC MRI data were collected on the same 3T Siemens TIM Trio Scanner and 32-channel head coil at the University of Pennsylvania for all participants. T1w images were acquired with a magnetization-prepared rapid acquisition gradient-echo (MPRAGE) sequence with the following parameters: TR = 1,810 ms, TE = 3.51 ms, TI = 1,100 ms, flip angle = 9 degrees, 160 slices, and voxel resolution = 0.94 × 0.94 × 1 mm. The single shell diffusion sequence consisted of b-value = 1,000 s/mm^2^ in 64 directions with 7 interspersed scans with b = 0 s/mm^2^. All 71 volumes were acquired in the anterior-posterior direction and were split between two runs. Diffusion scans were acquired with the following parameters: TR = 8,100 ms, TE = 82 ms, and voxel resolution = 1.875 × 1.875 × 2 mm. Furthermore, a map of the main magnetic field was acquired with a double-echo, gradient-recalled echo (GRE) sequence for susceptibility distortion correction of the diffusion data. The following parameters were used for field map acquisition: TR = 1,000 ms, TE = 2.69 and 5.27 ms, flip angle = 60 degrees, 44 slices, and voxel resolution = 3.75 × 3.75 × 4 mm.

HCP-D MRI data were collected at four sites on 3T Siemens Prisma scanners with 32-channel head coils as described above. T1w images were acquired using a 3D multi-echo MPRAGE sequence with an in-plane acceleration factor of 2 and the following parameters: TR = 2,500 ms, TE = 1.8, 3.6, 5.4, and 7.2 ms, TI = 1,000 ms, flip angle = 8 degrees, 208 slices, and voxel resolution = 0.8 mm isotropic. Multi-shell diffusion scans were acquired across four runs with b = 1,500 and 3,000 s/mm^2^ and a multiband factor of four. A total of 398 volumes were acquired, with 92-93 directions for each shell (370 directions total) and 28 b = 0 volumes. Two acquisitions with opposite phase encoding directions (anterior-posterior and posterior-anterior) were completed for each of the 185 unique directions. Diffusion scans used the following parameters: TR = 3,230 ms, TE = 89 ms, and voxel resolution = 1.5 mm isotropic. Reverse phase encoding EPI-based fieldmaps were acquired with the following parameters: TR = 8,000 ms, TE = 66 ms, flip angle = 90 degrees, 72 slices, and voxel resolution = 2.0 × 2.0 × 2.0 mm.

HBN MRI data were acquired on 3T MRI scanners at three different sites. At each site, the following scanners were used: 3T Siemens Tim Trio scanner at Rutgers University Brain Imaging Center and 3T Siemens Prisma Scanners at CitiGroup Cornell Brain Imaging Center and the CUNY Advanced Science Research Center. Participants scanned at Staten Island Flagship Research Center were not included in this study due to the use of a 1.5T Siemens Avanto scanner. T1w images were acquired with an MPRAGE sequence with the following parameters: TR = 2,500 ms, TE = 3.15 ms, TI = 1,060 ms, flip angle = 8 degrees, 224 slices, and voxel resolution = 0.8 mm isotropic. Multi-shell diffusion scans were acquired with a multiband factor of three with b = 1,000 and 2,000 s/mm^2^ in the anterior-posterior phase encoding direction. For each shell, 64 directions were acquired for 128 directions total. One b = 0 volume was acquired. The following parameters were used for the diffusion acquisition: TR = 3,320 ms, TE = 100.2 ms, and voxel resolution = 1.8 mm isotropic. A reverse phase encoding b = 0 was additionally acquired for use as an EPI-based field map in susceptibility distortion correction.

#### Diffusion MRI preprocessing

Preprocessing of diffusion scans, fieldmaps, and T1-weighted images, used QSIPrep^70^, an integrative pipeline for processing diffusion-weighted MRI data and uses the software tools described below. QSIPrep version 0.14.2 was used in PNC and HBN and version 0.16.1 in HCP-D, which included the following respective internal software versions: Nipype 1.6.1 and 1.8.5, Nilearn 0.8.0 and 0.9.2, ANTs 2.3.1 and 2.4.0, and FSL 6.0.3 and 6.0.5. The same preprocessing steps were applied to all datasets as described below.

The T1-weighted (T1w) image was corrected for intensity non-uniformity (INU) using N4BiasFieldCorrection^71^, and used as T1w-reference throughout the workflow. The T1w-reference was then skull-stripped using antsBrainExtraction (ANTs), using OASIS as the target template. Spatial normalization to the ICBM 152 Nonlinear Asymmetrical template version 2009c^72^ was performed through nonlinear registration with antsRegistration,^73^ using brain-extracted versions of both T1w volume and template. Brain tissue segmentation of cerebrospinal fluid, white-matter, and gray-matter was performed on the brain-extracted T1w using FAST (FSL)^74^. For PNC (2 runs) and HCP-D (4 runs), individual preprocessing steps were applied to each diffusion run and then subsequently concatenated; HBN data was all acquired in a single run. Any images with a b-value < 100 s/mm² were treated as a *b* = 0 image. MP-PCA denoising as implemented in MRtrix3’s dwidenoise^75^ was applied with a 5-voxel window. After MP-PCA, Gibbs unringing was performed using MRtrix3’s mrdegibbs^76^. Following unringing, B1 field inhomogeneity was corrected using dwibiascorrect from MRtrix3 with the N4 algorithm^71^. After B1 bias correction, the mean intensity of the diffusion-weighted series was adjusted such that the mean intensity of the b=0 images matched across separate runs. FSL’s eddy was used for head motion correction and Eddy current correction^77^. Eddy was configured with a q-space smoothing factor of 10, a total of 5 iterations, and 1000 voxels used to estimate hyperparameters. A linear first level model and a linear second level model were used to characterize Eddy current-related spatial distortion. q-space coordinates were forcefully assigned to shells. Field offset was attempted to be separated from subject movement. Shells were aligned post-eddy. Eddy’s outlier replacement was used^78^. Data were grouped by slice, only including values from slices determined to contain at least 250 intracerebral voxels. Groups deviating by more than 4 standard deviations from the prediction had their data replaced with imputed values.

Different approaches were used to correct for susceptibility artifacts as different versions of fieldmaps were acquired for PNC, HCP-D, and HBN. In the PNC, a B0 map using a phase-difference image and a magnitude image from the GRE fieldmap acquisition were created to assess susceptibility distortion correction. In HCP-D and HBN, reverse phase-encoding EPI-based fieldmaps were collected, resulting in pairs of images with distortions going in opposite directions. Here, b=0 reference images with reversed phase encoding directions were used along with an equal number of b=0 images extracted from the diffusion scans. From these pairs the susceptibility-induced off-resonance field was estimated^79^. The fieldmaps were ultimately incorporated into the Eddy current and head motion correction interpolation. Final interpolation for all datasets was performed using the Jacobian modulation method. As part of preprocessing, the diffusion data were resampled to AC-PC to be in alignment with the T1w image while retaining the input data resolution.

#### Additional structural MRI (sMRI) preprocessing

In addition to QSIPrep’s anatomical pipeline, additional processing of the T1-weighted image was required to generate the white and pial surfaces required for tract-to-cortex mapping (described below). Specifically, T1w images from all datasets were additionally processed with sMRIPrep 0.7.1 (as part of fMRIPrep 20.2.3^80^). T1w images underwent correction for intensity non-uniformity with N4BiasFieldCorrection from ANTs 2.3.3^71,73^, skull-stripping with a Nipype 1.6.1 implementation of ANTs brain extraction workflow, and brain tissue segmentation with fast FSL 5.0.9^74^. Cortical surfaces were then reconstructed using FreeSurfer 6.0.1^81^.

#### Reproducible image reconstruction

To ensure reproducibility and scalability of the reconstruction workflows described below (tractography and tractometry), we used BIDS App Bootstrap 0.0.8 (BABS; https://github.com/PennLINC/babs/releases/tag/0.0.8), a generalizable Python package for reproducible image processing^82^. BABS relies on DataLad^83^ and the FAIRly big framework^84^ to track a full audit trail of data processing of Brain Imaging Data Structure (BIDS)^85^ datasets and apps. BABS also generates code for data processing based on users’ customization. We used BABS to process PNC, HCP-D, and HBN through the reconstruction workflows in QSIPrep as described below.

#### Diffusion MRI reconstruction and tractography

We used custom reconstruction workflows in QSIPrep 0.22.0^70^ for dMRI reconstruction and tractography. Custom reconstruction json files may be found in the Github repository under “Code Availability”. This workflow first estimated fiber orientation distributions (FOD) of specific fiber populations using constrained spherical deconvolution (CSD). FODs served as the basis of biologically interpretable streamline tractography^86,87^. CSD overcomes limitations with crossing fibers and partial voluming present with diffusion tensor-based tractography (DTI)^86,88,89^. White matter tracts were reconstructed using multi-shell multi-tissue CSD tractography^89^ in HCP-D and HBN and single-shell 3-tissue CSD^90^ in PNC through the MRtrix3 software package^91^. Streamlines were mapped to corresponding FOD lobes. To refine endpoints of dMRI streamlines, Hybrid Surface-Volume Segmentation (HSVS) provided tissue interface localization information to Anatomically-Constrained Tractography (ACT). This tractography approach constrains the propagation and termination of streamlines to biologically plausible regions based on tissue segmentation of an anatomical T1-weighted image^92,93^. However, because utilizing ACT in HBN led to the reconstruction failure of over 100 participants, we did not apply ACT in this dataset. We conducted sensitivity analyses for HBN with and without ACT to confirm that this dataset-specific methodological variation did not significantly impact downstream results (**Figure S7**). We note that because HBN is a higher noise dataset, this led to generally greater challenges with tractography and tract segmentation (method described below) compared to PNC and HCP-D.

We employed two additional reconstruction workflows: AMICO^94^ to estimate the neurite orientation dispersion and density imaging (NODDI) model^29^ and TORTOISE^30^ to calculate mean apparent propagator (MAP) MRI fits^31^. NODDI and MAP-MRI leverage differential tissue responses from multiple *b*-values to model the cellular environment^22^. NODDI, which is a tissue-based model, estimates the directional distribution of axons and dendrites, or neurites, and matches diffusion patterns to this distribution. This model also estimates intracellular, extracellular, and isotropic volume fractions; we specifically examined the intracellular volume fraction (ICVF) from NODDI, a marker of neurite density^95^. MAP-MRI is a signal-based model, which models water molecule displacement without *a priori* assumptions about the tissue environment and directly estimates properties of the mean apparent propagator^22^. MAP-MRI models diffusion signal as a linear combination of angular and radial basis functions and estimates the 3D diffusion propagator and angular diffusion orientation distribution function^31^.

We specifically examined the return-to-origin probability (RTOP), or the probability of water molecules undergoing no net displacement in any direction^22^.

#### Tractometry analysis

We defined canonical white matter tracts using Automated Fiber Quantification in Python (pyAFQ).^96^ This work used pyAFQ 2.0 (https://github.com/tractometry/pyAFQ/releases/tag/2.0). As detailed in Kruper et al. 2024^97^, pyAFQ segmented white matter tracts in subject space based on inclusion and exclusion regions of interest^21,98^. Tract segmentation was performed on MRtrix3 tractography outputs from the QSIPrep reconstruction workflow described above. Of the 28 tracts delineated by pyAFQ, we excluded bilateral tracts that did not have two distinct cortical endpoints, including the cingulum bundle and anterior thalamic radiation. We included corticospinal tract as an exemplar tract that does not have two distinct cortical endpoints for sensitivity analyses.

To analyze microstructural properties along individual tracts, we computed tract profiles for each tract. First, we used the DTI model, implemented using DIPY, to estimate voxel-wise tensor-derived measures. Of note, the diffusion tensor model does not account for the non-gaussianity of water diffusion that occurs at higher b-values^22,99^. Because the multishell scans in HCP-D (b = 0, 1500, 3000) and HBN (b = 0, 1000, 2000) included high b-value shells, we excluded b-values > 1500. This value was chosen to maximize accuracy of DTI modeling while still leveraging multiple shells.

Next, we examined mean diffusivity along each tract, which may be more sensitive to developmental change and exhibits more widespread changes than fractional anisotropy in this age window^8,22,23^. Mean diffusivity was calculated at 100 equidistant nodes along each streamline in each tract. Tract profiles of mean diffusivity were calculated as the sum of each streamline’s mean diffusivity value at a given node, inversely weighted by the Mahalanobis distance of that node from the core location of the tract^21^. In addition to using anatomically-constrained tractography before tract segmentation (described above), the five end-most nodes on each end of each tract were excluded to minimize partial volume effects^21^.

To examine microstructural measures derived from multi-shell data (i.e. ICVF from NODDI and RTOP from MAP-MRI) along each tract, we employed the same approach described above using DIPY to generate tract profiles on white matter tracts segmented from pyAFQ.

#### Harmonization of tract profiles data

Multi-site tract profiles data underwent Correcting Covariance Batch Effects (CovBat) within each dataset where diffusion data were collected on multiple scanners (HCP-D and HBN)^100–102^. This scanner harmonization approach mitigates covariance-related batch effects while allowing us to account for non-linear age effects of interest. Specifically, we modeled age as a smooth term using a generalized additive model in both the initial mean-correction and the covariance-correction stages, similar to ComBat-GAM^103,104^. Sex and in-scanner motion were included as additional linear covariates. CovBat was implemented using the *CombatFamily* package (version 0.2.1) in R.

#### Tract to cortex mapping

We developed a workflow that allowed us to map tracts to their respective cortical endpoints by combining toolkits (e.g., pyAFQ, MRTrix, FSL, Freesurfer, Nilearn, sMRIPrep, and Connectome Workbench). This workflow overcame technical challenges of integrating data from different imaging modalities and registering cortical surfaces to volumetric white matter tracts while maximizing accuracy in defining cortical endpoints. With the resulting tract-to-cortex maps, we could relate WM development to each tract’s cortical endpoints.

To create tract-to-cortex probability maps for each dataset, we first transformed each subject’s FreeSurfer white matter surface to native AC-PC alignment to match the space of the subject’s QSIPrep T1-weighted image using FSL 6.0.4 and sMRIPrep 0.12.2. Next, we identified cortical endpoints for each tract. We binarized subject-level tract density images (TDIs) for each tract using ‘tckmap’ from MRTrix 3.0.4, creating maps that indicate whether a given tract was present or absent in each white matter voxel. Using the ‘vol_to_surf’ function in Nilearn 0.10.3, we sampled binarized TDI values 1.5 millimeters below the gray matter/white matter boundary, as defined by the FreeSurfer white matter surface. This depth was chosen by first sampling different depths below the gray matter/white matter boundary in the discovery dataset, PNC (0.5, 1, 1.5, and 2 mm). Using tissue probability maps generated by QSIPrep, we then determined that a depth of 1.5 mm yielded the lowest probability of sampling gray matter and the highest probability of sampling white matter. A greater depth of 2 mm sampled into the opposite cortical ribbon, whereas smaller depths may not have reached deep enough into white matter.

Subject-level surface maps of binarized TDI values were mapped to the fsLR surface using Connectome Workbench 1.4.2. These binarized fsLR surfaces were then averaged across subjects to create tract-to-cortex probability maps for each tract. These maps were then parcellated using the HCP-MMP atlas. For each dataset, each tract-specific map represents the proportion of participants with a tract termination at each HCP-MMP region. Cortical regions where at least 30% of subjects exhibited a non-zero TDI value were defined as cortical endpoints for most tracts. This threshold was selected to maximize accuracy of cortical endpoints based on known anatomy of tracts (**Figure S8**). For the inferior fronto-occipital fasciculus (IFOF), a 5% threshold was applied due to technical challenges in tract segmentation with anatomically-constrained tractography. Specifically, the IFOF must traverse a narrow bottleneck in the temporal cortex, making successful segmentation more challenging.

### QUANTIFICATION AND STATISTICAL ANALYSIS

#### Developmental models

To model linear and non-linear associations between age and mean diffusivity along tracts, generalized additive models (GAM) were fitted using the *mgcv* package (version 1.8.39) in R 4.1.2^105–109^. Separate GAMs were fit for each node within each tract, with mean diffusivity as the dependent variable, age as a smooth term, and sex and in-scanner motion as linear covariates.

As in previous work^4,68^, age was modeled using unpenalized thin plate regression splines as the smooth term basis, with a maximum basis complexity (*k*) of 3 to avoid overfitting, as model fits were generally not complex and required a smaller number of knots. The age spline represents the developmental fit of mean diffusivity for each node. For each node-wise GAM, we computed several developmental measures. The magnitude of the age effect was quantified by the absolute difference in adjusted *R*^2^ (Δ*R*^2^adj) between a full model and reduced model with no age term. The significance of the association between the mean diffusivity and age was assessed using analysis of variance (ANOVA) to compare the full and reduced models. Multiple comparisons were controlled for with false discovery rate (FDR) correction; *Q*<0.05. To determine age-specific rates of developmental change for each node, we used the ‘derivatives’ function in the *gratia* package (version 0.7.0). We computed the first derivative of the node-wise age smooth function using finite differences and calculated a simultaneous 95% confidence interval for the first derivative. Under the assumption that mean diffusivity decreases monotonically in this age window, the age of maturation was characterized as the first age at which this interval included zero, indicating that the rate of developmental change became non-significant. If no such age was identified, suggesting that the node matured beyond the window studied, the age of maturation for the node was set to the maximal age in the sample (for example, 23 years old for PNC). Of note, because deep tract regions exhibit very small magnitudes of age-related change, the age of maturation is not directly comparable to that of superficial tract regions, which demonstrated substantially larger developmental effects. Lastly, to examine change in mean diffusivity over age, we generated fitted values of mean diffusivity from the GAM at 200 timepoints between the minimum and maximum age of each study using the ‘fitted_values’ function in the *gratia* package. These same developmental models were used for tract profiles of multi-shell microstructural measures.

#### Characterizing development along entire lengths of tracts

##### Correspondence of along-tract developmental effects to a deep-to-superficial axis

To evaluate whether age effects were enriched in superficial tract regions compared to deep regions of each tract, we adapted recently developed tools for network enrichment significance testing (NEST)^110^ for highly spatially autocorrelated tract profile data. Of note, “superficial” does not refer to U-fibers, but rather distal regions along long-range WM tracts adjacent to the cortex. NEST creates a null distribution by permuting participant age while keeping brain data and other covariates fixed, preserving spatial structure in the data and controlling for Type I error rates. Using NEST, the age effect (Δ*R*^2^_adj_) was computed for deep (nodes 46-55) and superficial (nodes 5-9 and 90-94) tract regions, using a bin size of 5 nodes. To ensure robustness of our results to alternate specifications, we tested additional bin sizes (3, 7, and 10 nodes) for defining deep and superficial tract regions and found that bin size minimally influenced statistical significance (**Table S2**). NEST first ranked age effect magnitudes of each tract region, and then calculated an enrichment score to assess whether age effects were enriched in superficial versus deep tract regions. The observed enrichment score was compared against the null distribution generated by 10,000 permutations of age to compute a conservative, permutation-based *p-*value that accounts for the high degree of spatial autocorrelation along tracts.

#### Characterizing superficial tract region development

We first mapped tracts to their respective cortical endpoints as described above. Next, to investigate how developmental patterns of superficial tract regions correspond to a tract’s cortical endpoints, we mapped developmental measures to each tract’s endpoints. The developmental measures at the five most superficial nodes that were retained (nodes 5–9 and 90–94; as noted above, nodes 0-4 and 95-99 were removed in our analyses to mitigate partial volume effects) were averaged. We then plotted the averaged developmental measures on the cortical surface at the respective cortical endpoints for each tract. Mean diffusivity developmental trajectories for each endpoint were computed by averaging the fitted values for the five most superficial nodes on each end.

We then assessed whether age effects significantly differed between homotopic endpoints of a tract within the corpus callosum, specifically the motor segment of the corpus callosum (callosum motor). We also examined whether age effects differed between endpoints of a long-range white matter tract that connects heterotopic regions – the IFOF. Mean diffusivity age effects, as characterized by Δ*R*^2^adj, were computed for the five most superficial nodes and an enrichment score was calculated. For callosum motor, the five most superficial nodes in the right hemisphere were compared to their counterparts in the left hemisphere. For the IFOF, we compared the five superficial nodes adjacent to the bilateral frontal endpoints to the five nodes adjacent to the bilateral occipital endpoints. To evaluate if associations between age and mean diffusivity were enriched in one cortical endpoint compared to the other, we used NEST as described previously.

##### Correspondence of developmental patterns of superficial tract regions to the cortical hierarchy

We sought to determine if development of superficial tract regions aligned with the cortical hierarchy as defined by the sensorimotor-association (S-A) axis. Cortical endpoints for each tract were thus mapped to the S-A axis of cortical hierarchy^3^. The S-A axis spans from unimodal sensorimotor cortices to transmodal association cortices and was derived by averaging multimodal brain maps, including anatomical hierarchy quantified by T1w/T2w ratio^111^, functional hierarchy^112^, evolutionary hierarchy^113^, allometric scaling^114^, aerobic glycolysis^115^, cerebral blood flow^116^, gene expression^117^, first principal component of NeuroSynth terms^118^, externopyramidization^119^, and cortical thickness^3^. The rank ordering of HCP-MMP regions along the S-A axis (https://github.com/PennLINC/S-A_ArchetypalAxis) was used for this study.

Cortical regions included in each tract’s endpoints were assigned HCP-MMP parcellated S-A axis ranks and were averaged for each end. This process yielded two mean S-A ranks per tract, one for each endpoint.

Correspondence to the S-A axis was first examined at the tract-level. Pearson correlations quantified the association between the ages of maturation for each cortical endpoint and the mean S-A axis rank of that endpoint’s constituent cortical regions. We tested for statistical significance using spin-based spatial permutation tests, or “spin tests”, which mitigate distance-dependent spatial autocorrelation in cortical maps^120^. The spin test generates a null distribution by rotating spherical projections of one cortical feature map and computes a *p*-value (*p*_spin_) by comparing the empirically observed test statistic to the null. We adapted the ‘rotate_parcellation’ algorithm (https://github.com/frantisekvasa/rotate_parcellation)^121^ for tract-level endpoint data by spinning the S-A axis 10,000 times and recomputing an average spun S-A axis rank for each cortical endpoint. Of note, several cortical endpoints did not mature within the studied age window, resulting in a ceiling effect for ages of maturation. To address this issue, we supplemented our analyses by computing the Pearson correlation coefficients between average S-A rank and the age of maturation of endpoints that matured within the age range studied. Additionally, we used a two-sample, one-tailed *t*-test to investigate whether the S-A rank of still-developing endpoints was significantly greater than matured endpoints. We assessed statistical significance of this comparison using spin tests, as described above, with the *t*-value as the test statistic.

Next, we interrogated whether cortical endpoints at opposite ends of the cortical hierarchy show greater differences in age of maturation for each tract. We calculated two measures for each tract: the difference in age of maturation (ΔAge) and the difference in average S-A rank (ΔS-A rank). Given that one group of tracts exhibited relatively small ΔS-A rank and a second group exhibited large differences, we divided tracts into two groups (large and small ΔS-A rank) and implemented a one-tailed *t*-test to quantify whether the ΔAge was greater in tracts with large ΔS-A rank. Statistical significance was evaluated using a modified spin test in which differences in age of maturation were computed on spun age of maturation maps, and a null distribution of *t*-values was generated by conducting *t*-tests on null ΔAge values. The age of maturation maps used for the null distribution were created through the following steps. For each dataset, we aggregated the ages of maturation for superficial tract regions across all tracts and assigned the average age of maturation to the nearest cortical endpoint. This process yielded an average age of maturation map across all superficial tract regions for each dataset, with each HCP-MMP region that had a tract termination being assigned an average value. Of note, we utilized all endpoints, including ones that did not fully mature in the age window, to retain enough data points for this analysis.

Lastly, using the across-tract age of maturation maps described above for each dataset, Pearson correlations quantified the association between the aggregated age of maturation map and the S-A axis of each HCP-MMP region. To address ceiling effects caused by endpoints maturing beyond the studied age window, we repeated this analysis using Pearson correlation coefficients to evaluate the association between the age of maturation of matured regions and the S-A axis. We additionally performed a two-sample, one-tailed *t*-test to determine whether the S-A ranks of still-developing regions were significantly greater than those of matured regions. Statistical significance for these analyses was determined using spin tests.

## RESOURCE AVAILABILITY

### Lead Contact

Further information and requests regarding resources should be directed to and will be fulfilled by the lead contact, Theodore D. Satterthwaite.

### Materials Availability

This study did not generate new unique reagents.

### Data and Code Availability

This paper analyzes publicly available data from three datasets: the Philadelphia Neurodevelopmental Cohort, accessible from the Database of Genotypes and Phenotypes (phs000607.v3.p2) at https://www.ncbi.nlm.nih.gov/projects/gap/cgi-bin/study.cgi?study_id=phs000607.v3.p2; Human Connectome Project: Development is available for download through the NIMH Data Archive (https://nda.nih.gov/); and Healthy Brain Network is accessible through https://fcon1000.projects.nitrc.org/indi/cmihealthybrainnetwork/.

All analysis code is available at https://github.com/PennLINC/luo_wm_dev. A detailed guide to the code and procedures for replicating all analyses can be found at https://pennlinc.github.io/luo_wm_dev.

## ACKNOWLEDGEMENTS

This study was supported by grants from the National Institute of Health: R01MH120482 (T.D.S. and M.P.M.), R01MH113550 (T.D.S.), R01MH112847 (R.T.S. and T.D.S.), R01EB022573 (T.D.S.), RF1MH116920 (T.D.S.), R37MH125829 (T.D.S.), RF1MH121868 (A.R.), RF1MH121867 (A.R.), R01EB027585 (A.R.), RF1MH126699 (A.R.), R01MH119219 (R.E.G. and R.C.G.), R01MH123550 (R.T.S.), R01NS060910 (R.T.S.), R01MH123563 (R.T.S.), R00MH127293 (B.L.), 5T32MH019112-32 (A.S.K. & S.L.M.), 1L30MH131061-01 (A.S.K.), T32MH016804 (V.J.S.), R01MH120174 (D.RR.), R01MH132934 (A.A.B.), R01MH133843 (A.A.B.), U01MH109589 (D.M.B. & L.H.S), and T32GM007170 (F.H.). A.R. was additionally supported by a National Science Foundation grant (1934292) and by the Chan Zuckerberg Initiative’s Essential Open Source Software for Science program for software development. A.S.K. was additionally supported by a NARSAD Young Investigator Award from the Brain and Behavior Research Foundation. Additional support was provided by NIH U24NS130411, the AE Foundation, the Center for Artificial Intelligence and Data Science for Integrated Diagnostics at Penn, and the Penn/CHOP Lifespan Brain Institute.

## AUTHOR CONTRIBUTIONS

Conceptualization, A.C.L.; software, A.C.L., M.C., A.R., J.D.Y., S.L.M., M.J., S.M.W.; data curation, A.C.L., M.C., S.L.M.; resources, A.C.L., M.C., A.R., D.M.B., C.D., A.R.F., R.E.G., R.C.G., G.K., M.P.M, D.R.R., L.H.S., J.D.Y., T.D.S.; formal analysis, A.C.L.; supervision, T.D.S.; validation, S.L.M.; investigation, A.C.L.; visualization, A.C.L.; methodology, A.C.L.; writing – original draft, A.C.L.; writing – review & editing, S.L.M., V.J.S., A.A.B., J.B., D.M.B., D.S.B., C.D., A.R.F., J.G., R.E.G., R.C.G., F.H., M.J., G.K., A.S.K., B.L., A.P.M., M.P.M., D.R.R., G.S., R.T.S., L.H.S., S.M.W., J.D.Y., M.C., A.R., T.D.S.

## SUPPLEMENTAL INFORMATION

Document S1. Figures S1-S8. Tables S1 and S2.

