## Supplementary Information for "Two Axes of White Matter Development"

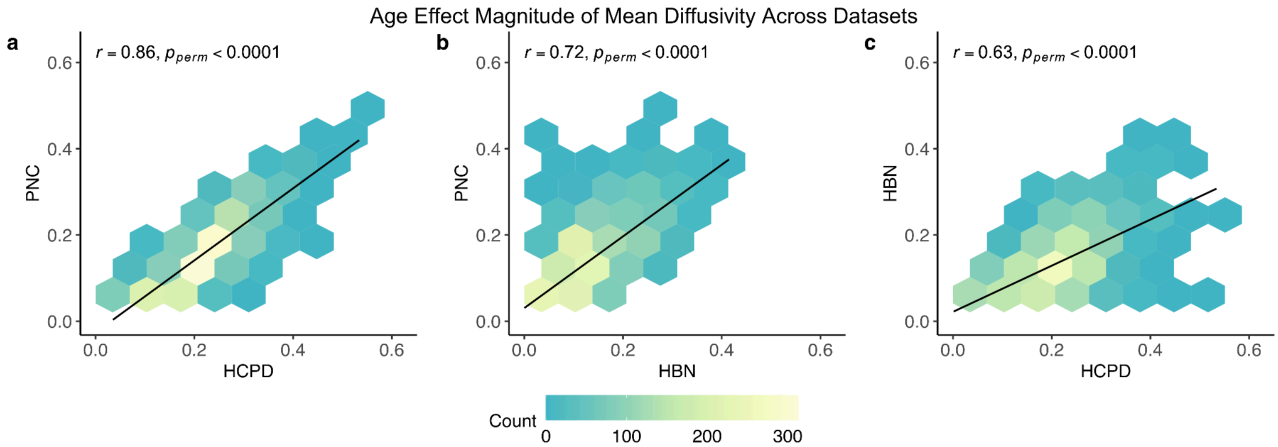


**Figure S1**. **Correspondence of mean diffusivity age effects between datasets. Related to Figures 1 and 3.** The magnitude of age-related change in mean diffusivity at each node (collapsed across all tracts) exhibits very high correspondence between datasets. The hexplots display the correspondence between **(a)** the Philadelphia Neurodevelopmental Cohort (PNC) and Human Connectome Project: Development (HCP-D) ($r$ = 0.86, $p_{\text{perm}} < 0.0001$), **(b)** PNC and Healthy Brain Network (HBN) ($r$ = 0.72, $p_{\text{perm}} < 0.0001$), and **(c)** HBN and HCP-D ($r$ = 0.63, $p_{\text{perm}} < 0.0001$). Pearson correlations were used to quantify the correlations between each pair of datasets. Statistical significance was determined by comparing the empirical correlation to a null distribution generated by shuffling the age effect across nodes 10,000 times.

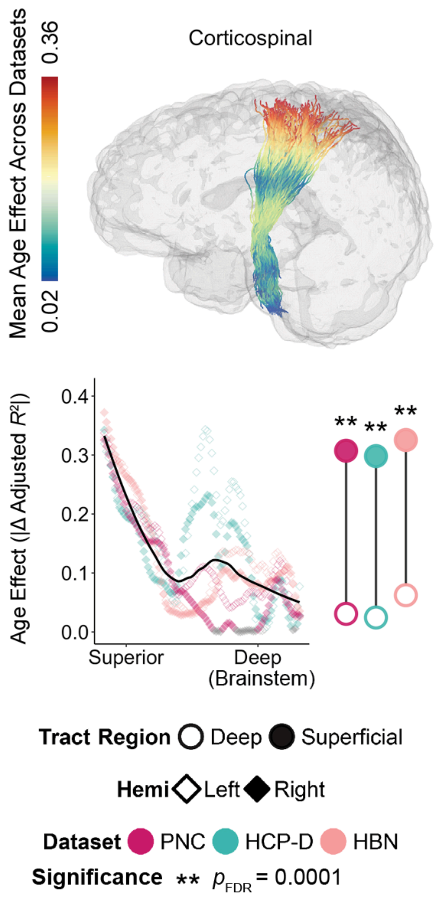


**Figure S2. The corticospinal tract exhibits a deep-to-superficial developmental patterning. Related to Figure 3.** The absolute value of the mean diffusivity age effect varies continuously along a projection tract, the corticospinal tract. The plot displays the magnitude of the age effect at 100 equidistant nodes that span the length of the tract. Data points are colored by dataset: the Philadelphia Neurodevelopmental Cohort (PNC), Human Connectome Project: Development (HCP-D), and Healthy Brain Network (HBN). Nodes without a significant age association are colored in gray ($Q_{FDR}$ > 0.05). Open and closed diamond shapes indicate left and right hemisphere tracts, respectively. The black line shows the overall trend, averaged across datasets and hemispheres. To the right of the age effect plot, average age effects for superficial (filled circle) and deep (open circle) tract regions are shown for each dataset. Significant differences between age effects in superficial (adjacent to the motor cortex) and deep (near the brainstem) tract regions were assessed using a network enrichment significance test. Stars denote significance levels following FDR correction.

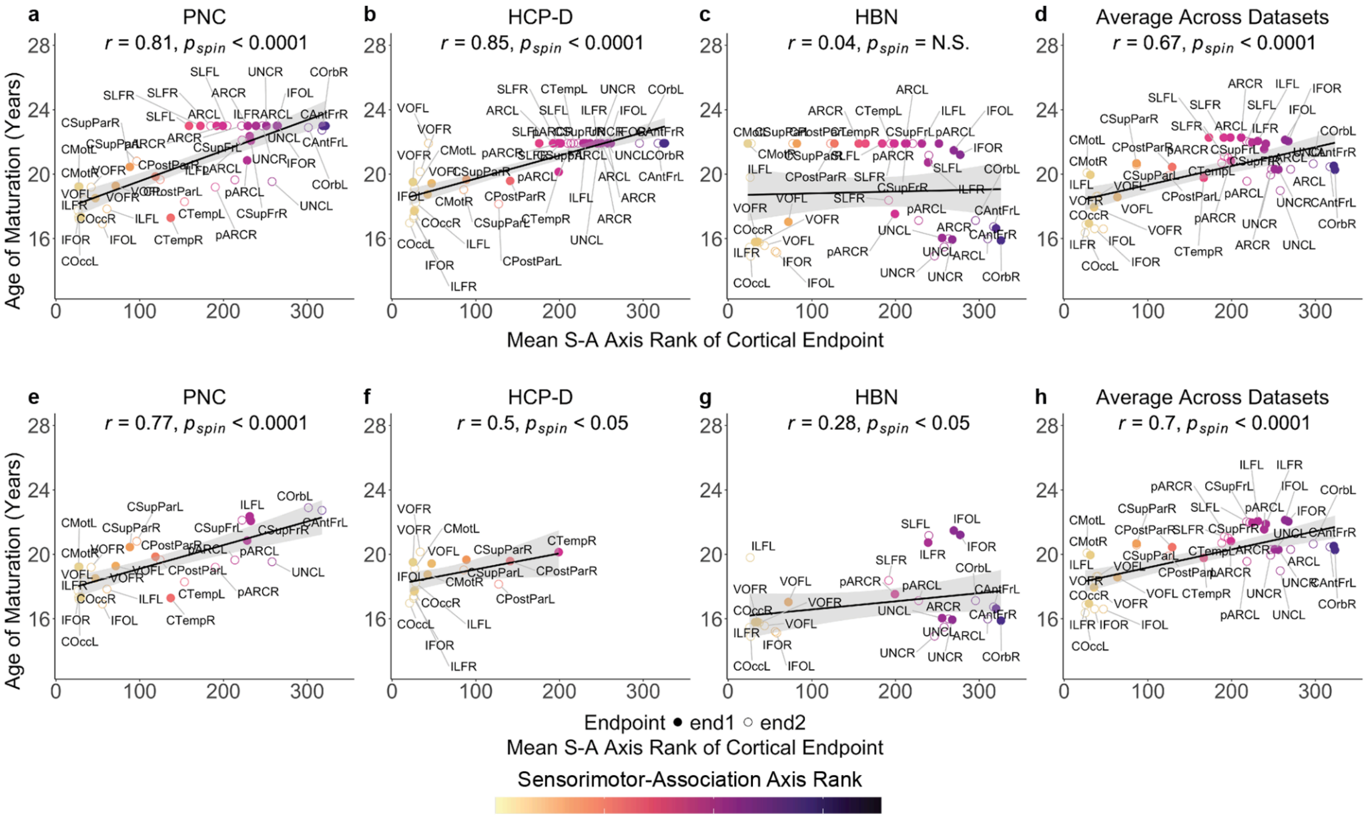


**Figure S3. Age of maturation for tract endpoints is associated with average sensorimotor-association axis rank of each endpoint's constituent cortical regions. Related to Figure 6.** We created tract-level maps of age of maturation by averaging the ages of maturation for the most superficial nodes within each tract on each end. Age of maturation in superficial tract regions is largely explained by each endpoint's mean sensorimotor-association (S-A) axis rank in **(a)** the PNC ($r$ = 0.81, $p_{spin}$ < 0.0001) and **(b)** HCP-D: $r$ = 0.85, $p_{spin}$ < 0.0001, but not in **(c)** HBN: $r$ = 0.04, $p_{spin}$ = N.S.). **(d)** When averaged across datasets, a significant effect remains ($r$ = 0.67, $p_{spin}$ < 0.0001). It should be noted that many endpoints did not reach maturation within the studied age window, creating a ceiling effect when examining the relationship between age of maturation and S-A rank. Excluding still-developing endpoints yields consistent results. After exclusion, age of maturation remains associated with S-A rank in **(e)** PNC ($r$ = 0.77, $p_{spin}$ < 0.0001), **(f)** HCP-D ($r$ = 0.5, $p_{spin}$ < 0.05), and **(g)** HBN ($r$ = 0.28, $p_{spin}$ < 0.05). **(h)** Averaged across datasets, a significant correlation is observed ($r$ = 0.7, $p_{spin}$ < 0.0001). Pearson correlations were used to quantify the association between the ages of maturation at each endpoint with the mean S-A rank of that endpoint's constituent cortical regions. Statistical significance was assessed using spin tests.

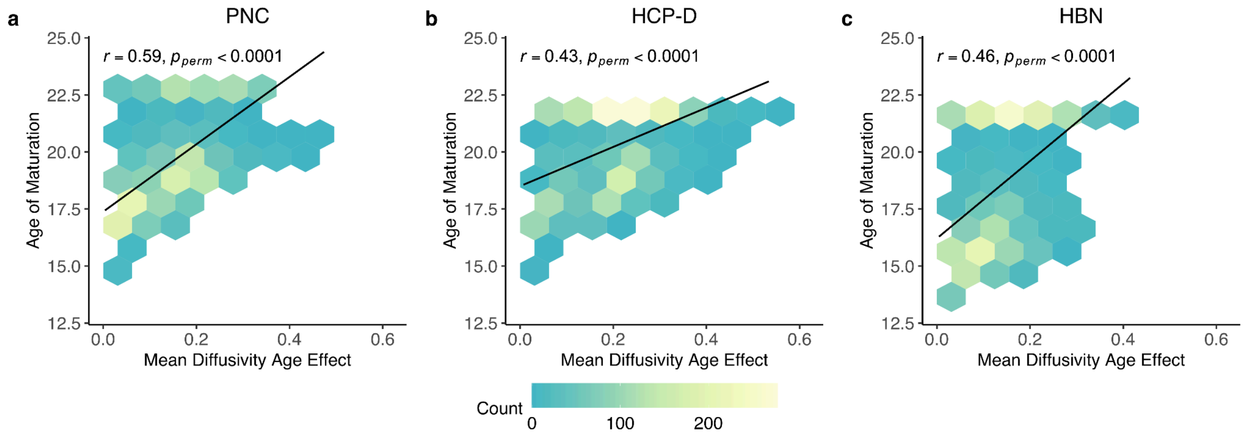


**Figure S4**. **Correspondence of mean diffusivity age effect and age of maturation. Related to Figure 6.** The magnitude of mean diffusivity age effects at each node (collapsed across all tracts) exhibits a moderate correspondence with age of maturation across datasets. The hexplots display the correlations in **(a)** the PNC ($r$ = 0.59, $p_{\mathrm{perm}} < 0.0001$), **(b)** HCP-D ($r$ = 0.43, $p_{\mathrm{perm}} < 0.0001$), and **(c)** HBN ($r$ = 0.46, $p_{\mathrm{perm}} < 0.0001$). While these two measures capture overlapping aspects of development, they also provide distinct developmental information from each other and along tracts. For instance, while the effect size of age is generally small in deep tract regions, the ages of maturation are quite variable. Some deep tracts tract regions that have a modest age effect nonetheless undergo small, protracted refinement throughout the age range studied. Age of maturation in deep tract regions thus captures the slowing of relatively small refinements. In contrast, superficial tract regions exhibit dramatic age-related changes in the studied age window. Thus, age of maturation in these superficial tract regions corresponds to the slowing of pronounced changes. Pearson correlations were used to quantify the correlations between each pair of datasets. Statistical significance was determined by comparing the empirical correlation to a null distribution generated by shuffling the age effect 10,000 times.

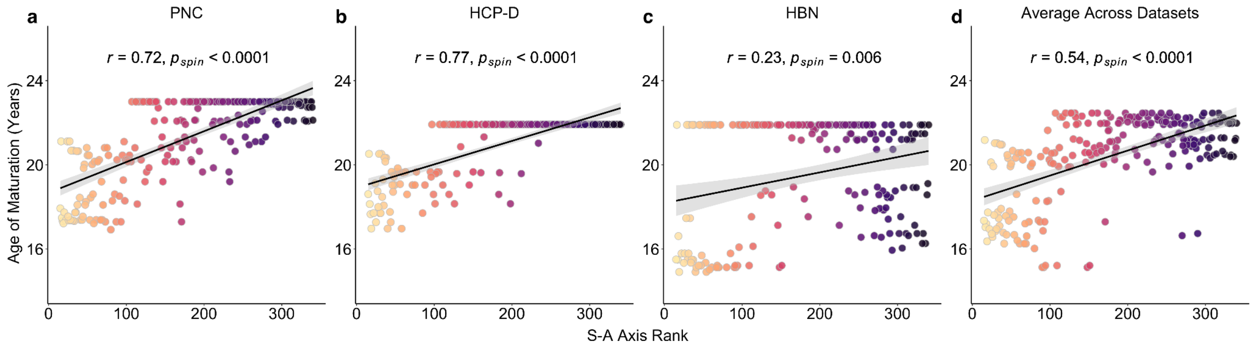


**Figure S5. Age of maturation across tracts is associated with the sensorimotor-association axis. Related to Figure 7.** Parcel-level cortical maps were created by averaging age of maturation maps across all superficial tract regions for each dataset, with each HCP-MMP region that had a tract termination being assigned an average age of maturation value. The age of maturation of all cortical regions that had a tract termination is associated with S-A axis rank in each dataset. Regions with older ages of maturation rank higher on the S-A axis in all three datasets: **(a)** PNC ($r = 0.72$, $p_{spin} < 0.0001$), **(b)** HCP-D (HCP-D: $r = 0.77$, $p_{spin} < 0.0001$), and **(c)** HBN ($r = 0.23$, $p_{spin} = 0.006$), as well as **(d)** across all datasets ($r = 0.54$, $p_{spin} < 0.0001$). Statistical significance of the Pearson correlation coefficients was assessed using region-based spin tests.

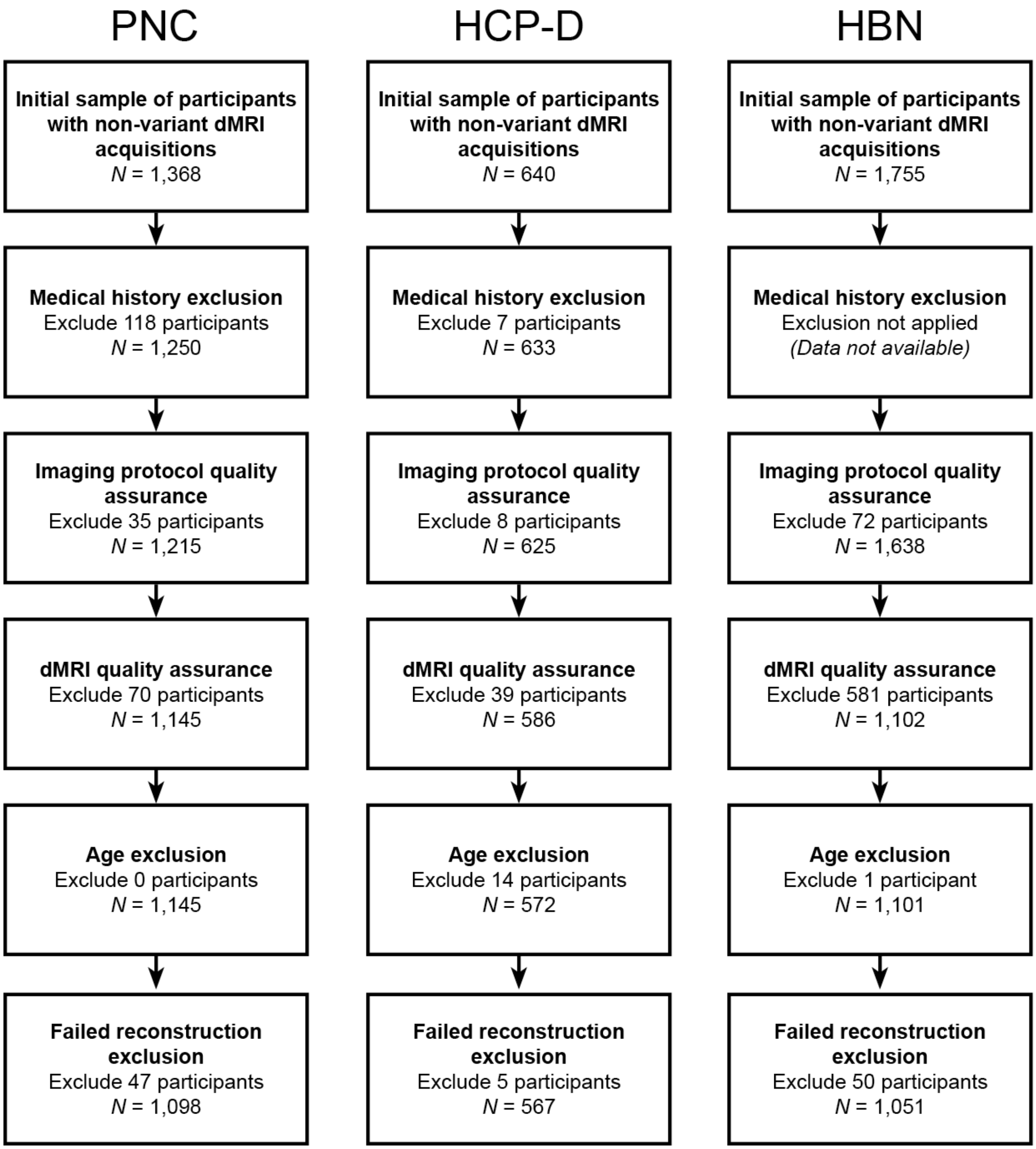


**Figure S6**. **Flow diagram depicting sample construction as well as inclusion and exclusion criteria. Related to STAR Methods.** Sample construction is shown for Philadelphia Neurodevelopmental Cohort (PNC), Human Connectome Project: Development (HCP-D), and Healthy Brain Network (HBN). Participants were excluded for variant acquisitions, medical conditions affecting brain function or gross neurological abnormalities, imaging protocol quality, diffusion MRI quality, age exclusion, and failed reconstruction.

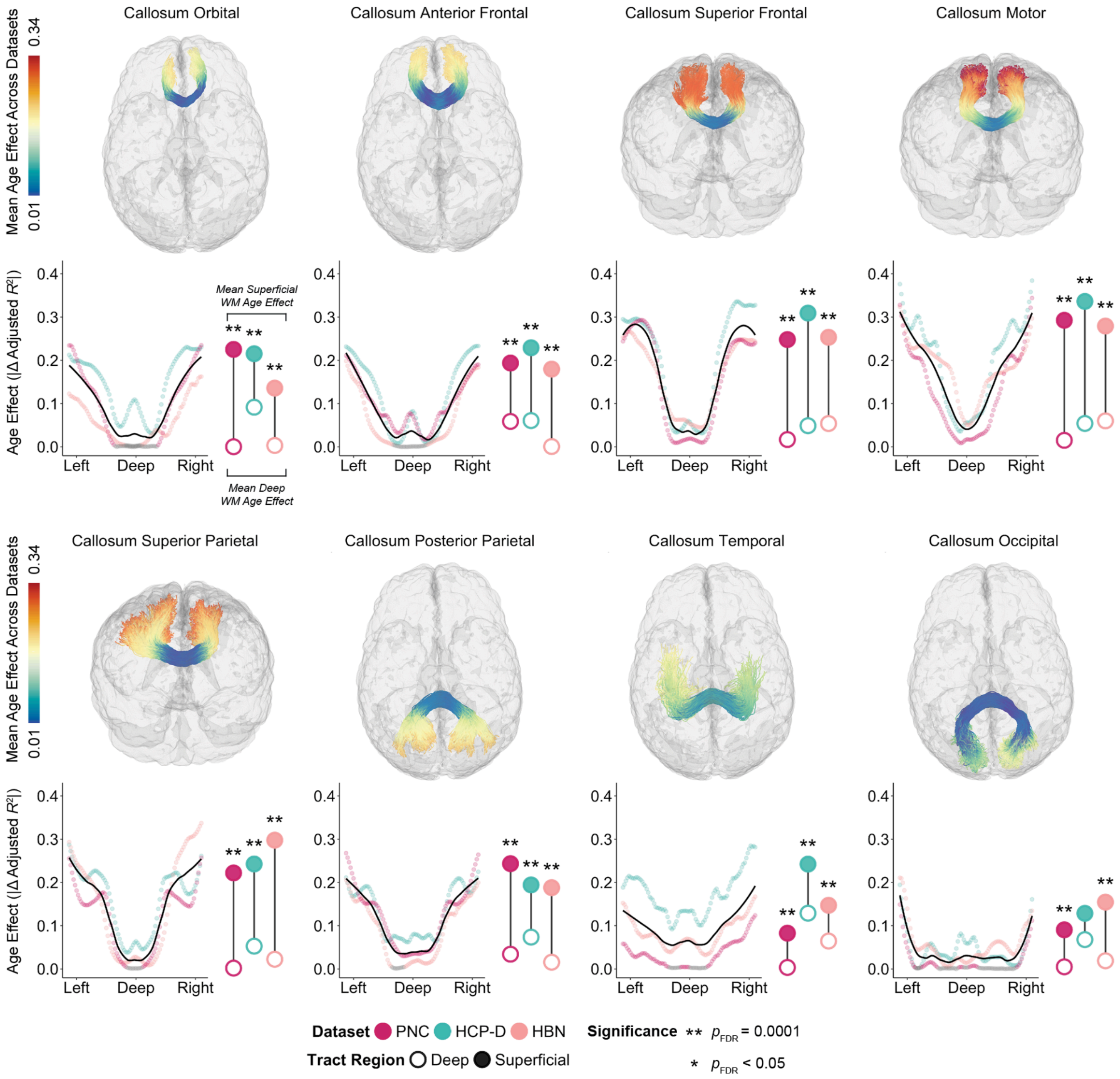


**Figure S7. Age effect of mean diffusivity in Healthy Brain Network processed with anatomically constrained tractography (ACT). Related to Figure 1 and STAR Methods.** The magnitudes of the age effect for mean diffusivity in Healthy Brain Network (HBN) processed with anatomically-constrained tractography (ACT) appear highly similar to that of HBN processed without ACT, as depicted in Figure 1.

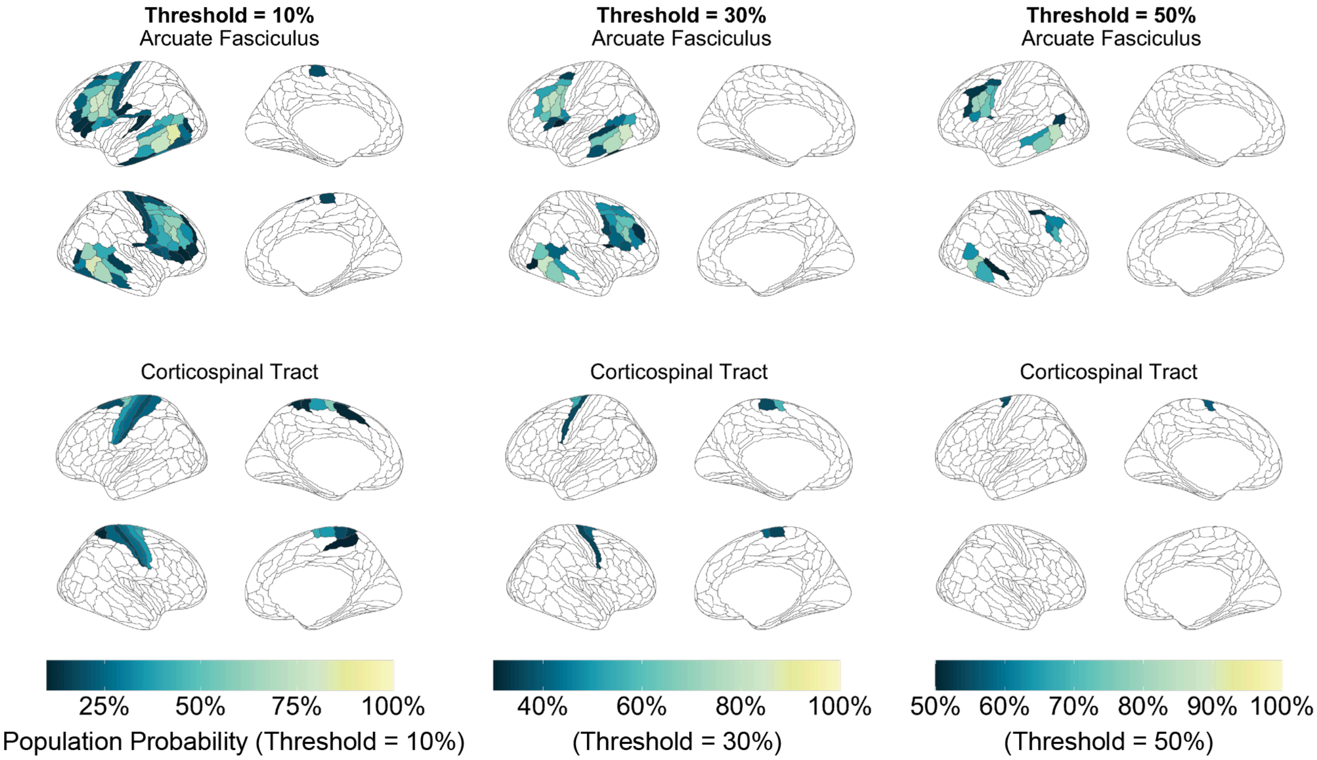


**Figure S8. Optimal threshold for determining cortical endpoints for tracts. Related to STAR Methods.** We computed tract-to-cortex probability maps for each tract, which were parcellated using the HCP-MMP atlas. The probability corresponds to the proportion of subjects exhibiting a termination at a given HCP-MMP region. Cortical regions that met the threshold probability were defined as cortical endpoints for each tract. Cortical endpoints for arcuate fasciculus and corticospinal tract are displayed for 10%, 30%, and 50%. A threshold of 30% yielded the most accurate cortical endpoints based on known anatomy of tracts. Tract-to-cortex probability maps from the PNC are shown.

| | Average Coefficient of Variation | | | | | |
|---|---|---|---|---|---|---|
| | PNC | | HCP-D | | HBN | |
| Tract | Deep | Superficial | Deep | Superficial | Deep | Superficial |
| L Arcuate | 4.26 | 4.11 | 3.81 | 3.92 | 7.95 | 5.31 |
| R Arcuate | 4.95 | 4.18 | 4.24 | 3.8 | 7.45 | 5.62 |
| Callosum Anterior Frontal | 5 | 3.57 | 4.35 | 3.56 | 8 | 4.94 |
| Callosum Motor | 12.13 | 4.65 | 8.93 | 3.82 | 11.09 | 7.06 |
| Callosum Occipital | 7.88 | 4.45 | 4.64 | 4.33 | 13.96 | 5.74 |
| Callosum Orbital | 7.76 | 4.1 | 4.44 | 3.95 | 12.29 | 6.59 |
| Callosum Posterior Parietal | 5.52 | 4.93 | 3.96 | 3.82 | 13.33 | 6.55 |
| Callosum Superior Frontal | 11.18 | 4.89 | 7.29 | 3.94 | 9.01 | 6.5 |
| Callosum Superior Parietal | 9.18 | 4.38 | 4.97 | 3.65 | 11.85 | 6.45 |
| Callosum Temporal | 10.26 | 6.18 | 5.8 | 4.11 | 18.64 | 12.47 |
| L Corticospinal | 3.19 | 11.09 | 3.26 | 6.35 | 7.47 | 9.05 |
| R Corticospinal | 3.44 | 11.14 | 2.92 | 6.73 | 7.19 | 9.2 |
| L Inferior Fronto.occipital | 9.18 | 4.65 | 5.92 | 4.35 | 7.79 | 5.63 |
| R Inferior Fronto.occipital | 5.69 | 5.23 | 5.64 | 4.49 | 7.36 | 5.55 |
| L Inferior Longitudinal | 4.9 | 4.66 | 4.11 | 3.97 | 6.93 | 6.69 |
| R Inferior Longitudinal | 5.77 | 4.69 | 4.14 | 3.84 | 7.22 | 6.05 |
| L Posterior Arcuate | 5.09 | 4.23 | 4.31 | 3.68 | 6.25 | 4.42 |
| R Posterior Arcuate | 5.3 | 4.63 | 4.18 | 3.7 | 6.4 | 5 |
| L Superior Longitudinal | 4.29 | 4.37 | 3.84 | 3.9 | 7.55 | 5.3 |
| R Superior Longitudinal | 4.54 | 4.38 | 3.51 | 3.83 | 7.44 | 5.46 |
| L Uncinate | 3.64 | 4.53 | 4.59 | 4.33 | 7.41 | 7.05 |
| R Uncinate | 3.47 | 4.36 | 3.51 | 4.17 | 6.85 | 6.42 |
| L Vertical Occipital | 5.78 | 4.96 | 4.09 | 4.79 | 6.05 | 5.64 |
| R Vertical Occipital | 5.82 | 4.56 | 4.29 | 4.31 | 5.87 | 5.66 |
| Average Across Tracts | 6.18 | 5.12 | 4.61 | 4.22 | 8.81 | 6.43 |

**Table S1. Average coefficient of variation in deep and superficial tract regions. Related to Figures 1 and 3.** The table displays the average coefficient of variation (CV) for deep and superficial tract regions for each tract in all three datasets. The average CV across all tracts is shown at the bottom of the table. Superficial tract regions of most tracts show lower CV than that in deep tract regions, suggesting that greater magnitudes of age effects in superficial regions are not driven by increased variability.

| | p-value | | | |
|---|---|---|---|---|
| Tract | Bin Size 3 | Bin Size 5 | Bin Size 7 | Bin Size 10 |
| Callosum Anterior Frontal | 0.00012 | 0.00012 | 0.00012 | 0.00012 |
| Callosum Motor | 0.00012 | 0.00012 | 0.00012 | 0.00012 |
| Callosum Occipital | 0.00012 | 0.00012 | 0.00012 | 0.00012 |
| Callosum Orbital | 0.00012 | 0.00012 | 0.00012 | 0.00012 |
| Callosum Posterior Parietal | 0.00012 | 0.00012 | 0.00012 | 0.00012 |
| Callosum Superior Frontal | 0.00012 | 0.00012 | 0.00012 | 0.00012 |
| Callosum Superior Parietal | 0.00012 | 0.00012 | 0.00012 | 0.00012 |
| Callosum Temporal | 0.00012 | 0.00012 | 0.00012 | 0.00012 |
| Arcuate | 0.00012 | 0.00012 | 0.00012 | 0.00012 |
| Inferior Fronto-occipital | 0.00012 | 0.00012 | 0.0084 | 0.0084 |
| Inferior Longitudinal | 0.39 | 0.23 | 0.27 | 0.27 |
| Posterior Arcuate | 0.00012 | 0.00012 | 0.00012 | 0.00012 |
| Superior Longitudinal | 0.00012 | 0.00012 | 0.00012 | 0.00012 |
| Uncinate | 0.16 | 0.14 | 0.12 | 0.095 |
| Vertical Occipital | 0.00012 | 0.00012 | 0.00012 | 0.00012 |

**Table S2. Bin sizes used for network enrichment significance testing (NEST) do not impact statistical significance. Related to STAR Methods.** Varying bin sizes used to define nodes in deep and superficial tract regions does not alter the statistical significance of permuted *p*-values of the mean diffusivity age effect generated by network enrichment significance testing. *P*-values comparing the age effect in deep versus superficial tract regions are shown. Representative results are displayed for the PNC, with similar findings observed in HCP-D and HBN.
